# VISTA-FISH: Video Imaging with Spatial-Temporal Analysis by Fluorescent In Situ Hybridization

**DOI:** 10.1101/2025.11.03.686351

**Authors:** Kun H. Lee, Mingjia Yao, Javid Ghaemmaghami, Cynthia J. DeLong, Neil Zhao, Jaimee Moline, Jonathan Sexton, Sami J. Barmada, Joshua D. Welch

## Abstract

Performing live-cell microscopy and high-dimensional gene expression measurements on the same cells is crucial for unraveling the molecular mechanisms underlying complex temporal phenotypes, yet this remains challenging using traditional approaches. To address these limitations, we developed Video Imaging with Spatial-Temporal Analysis by FISH (VISTA-FISH), a technique that matches live-cell recordings of cultured cells with the expression of thousands of genes in the same cell at the end of the video. Moreover, VISTA-FISH can simultaneously detect CRISPR guide RNAs in pooled screens, enabling investigation into the fundamental underpinnings of cellular activity, organelle transport, and other pivotal cellular functions. Using VISTA-FISH, we measured gene expression and neuron activity in the same differentiating neurons. This combined single-cell transcriptomic and video data allowed us to link differentiation stage, subcellular transcript localization, and cell subtype with neuron activity. Using these measurements, we built a model to predict activity from expression. Finally, we performed a pooled CRISPR interference screen with live-cell lysosome imaging, identifying alterations in lysosome movement and morphology as well as gene expression in perturbed neurons. Collectively, these studies position VISTA-FISH as a powerful new tool for elucidating the molecular mechanisms underlying neuron activity, organelle trafficking, and other complex temporal phenotypes.

## Introduction

Deciphering the molecular mechanisms that underlie complex cellular phenotypes remains a fundamental challenge in biology. Current methods can typically measure cellular changes over time or high-dimensional molecular states at a single time point, but not both. High-throughput single-cell RNA sequencing (scRNA-seq) enables transcriptome-wide profiling at single-cell resolution, characterizing novel cell types and transcriptional states^1–3^. However, scRNA-seq requires tissue dissociation and kills the cell, resulting in the loss of spatial information and making it challenging to link the gene expression profile of the cell with other cellular properties^4,5^. Conversely, live-cell microscopy measures cell shape and dynamic behaviors such as neuron firing or organelle movement but cannot measure many genes at once, making it difficult to link gene expression to temporal and morphological phenotypes. Consequently, most existing methods provide either molecular or temporal insights into cell biology, but rarely both in the same context.

Spatial transcriptomics (ST) preserves the physical organization of cells within intact tissue, enabling the study of cellular gene expression within the native microenvironment.

Recent advances in ST offer high-throughput gene expression measurement using sequential imaging or spatially barcoding sequencing^4,6^. Each category offers distinct benefits in spatial resolution, transcriptome coverage, and technical complexity, making them suited for different biological applications. For example, imaging-based ST provides subcellular resolution but is limited to targeted gene panels, while sequencing-based approaches offer unbiased whole-transcriptome coverage at lower spatial resolution.

Measuring gene expression while retaining spatial position opens an exciting new possibility: measuring temporal properties of cells using live-cell microscopy, then linking these images with cellular gene expression in the same cells. Because cells stay in the same place during ST measurement, the spatial position of each cell serves as a unique feature that can identify the same cell in live-cell videos and ST results. Previous studies captured images of fixed cells after performing ST, but the possibility of linking ST with live-cell imaging has remained largely unexplored. Investigators can transfer frozen or fixed sections onto the surface used for ST, but this complicates the process of identifying the same cells from living tissue in the spatial transcriptomic assay. However, culturing cells directly on the surface used for spatial transcriptomics makes it possible to match cells from live-cell microscopy and spatial transcriptomics, because the cells remain in the same locations for both measurements.

In this study, we introduce Video Imaging with Spatial-Temporal Analysis by Fluorescent In Situ Hybridization (VISTA-FISH), a new experimental approach that records videos from thousands of living cells at once, then uses spatial transcriptomics to measure expressed genes in individual cells at the end of the video. We accomplish this by maintaining cells directly on a surface compatible with spatial transcriptomics, allowing us to perform longitudinal imaging of living cells before spatial transcriptomics. These videos are then linked with subsequent single-cell gene expression measurements using spatial registration. We further extend VISTA-FISH to capture CRISPR guide RNAs, allowing us to record videos of living cells that have been genetically perturbed, measure their gene expression at the end of the video, and retrospectively link the perturbation that each cell received to its video and gene expression. We developed VISTA-FISH using the 10X Genomics Xenium platform, an imaging-based ST system that uses sequential FISH to detect transcripts from up to 5,000 genes with 30 nm resolution.

We used VISTA-FISH to simultaneously measure neural activity and gene expression in induced pluripotent stem cell (iPSC)-derived neural cells. We also performed a pooled CRISPR interference screen on iPSC-derived neurons and assessed resulting changes in lysosomal dynamics and cell morphology. Although we focused on neurons in this study, VISTA-FISH offers a flexible framework that integrates live-cell imaging, spatial transcriptomics, CRISPR perturbation, and immunostaining, and is broadly applicable to diverse cell types. By bridging molecular states with dynamic phenotypes, VISTA-FISH represents a powerful approach for studying molecular mechanisms underlying complex cellular phenotypes from a range of domains, including neuroscience, developmental biology, and immunology.

## Results

### VISTA-FISH records a video and gene expression from the same cells

ST technologies provide a unique opportunity to link molecular and imaging data, because the molecular measurement does not require removing the cells from their spatial positions.

Conventional ST approaches utilize thin tissue sections–cut from tissue containing cells that are already dead–adhered to surfaces compatible with ST. This approach allows linking static images of H&E or antibody staining with gene expression, but lacks the ability to connect phenotypes observed in live cells to their individual transcriptomic signature. In contrast, directly culturing living cells on a spatial transcriptomic slide should allow us to couple temporal, live-cell videos with gene expression. Methods that directly associate gene expression with temporal phenotypes at single-cell resolution in this way hold great promise for the discovery of molecular pathways that regulate complex aspects of cell biology.

Toward this goal, we developed Video Imaging with Spatial-Temporal Analysis by Fluorescent In Situ Hybridization (VISTA-FISH, **Fig. 1**). The key idea of VISTA-FISH is to culture cells directly on slides compatible with ST, allowing for live-cell imaging and ST profiling. After ST measurement, the cells can undergo additional rounds of immunostaining for proteins and other markers of interest. Finally, the same cells can be identified in live-cell images, ST images, and immunostained images by registering their spatial positions. Thus, VISTA-FISH yields a live-cell video for each cell, along with subcellular-resolution transcript locations and staining images for the same cell at the end of the video.

**Figure 1.**
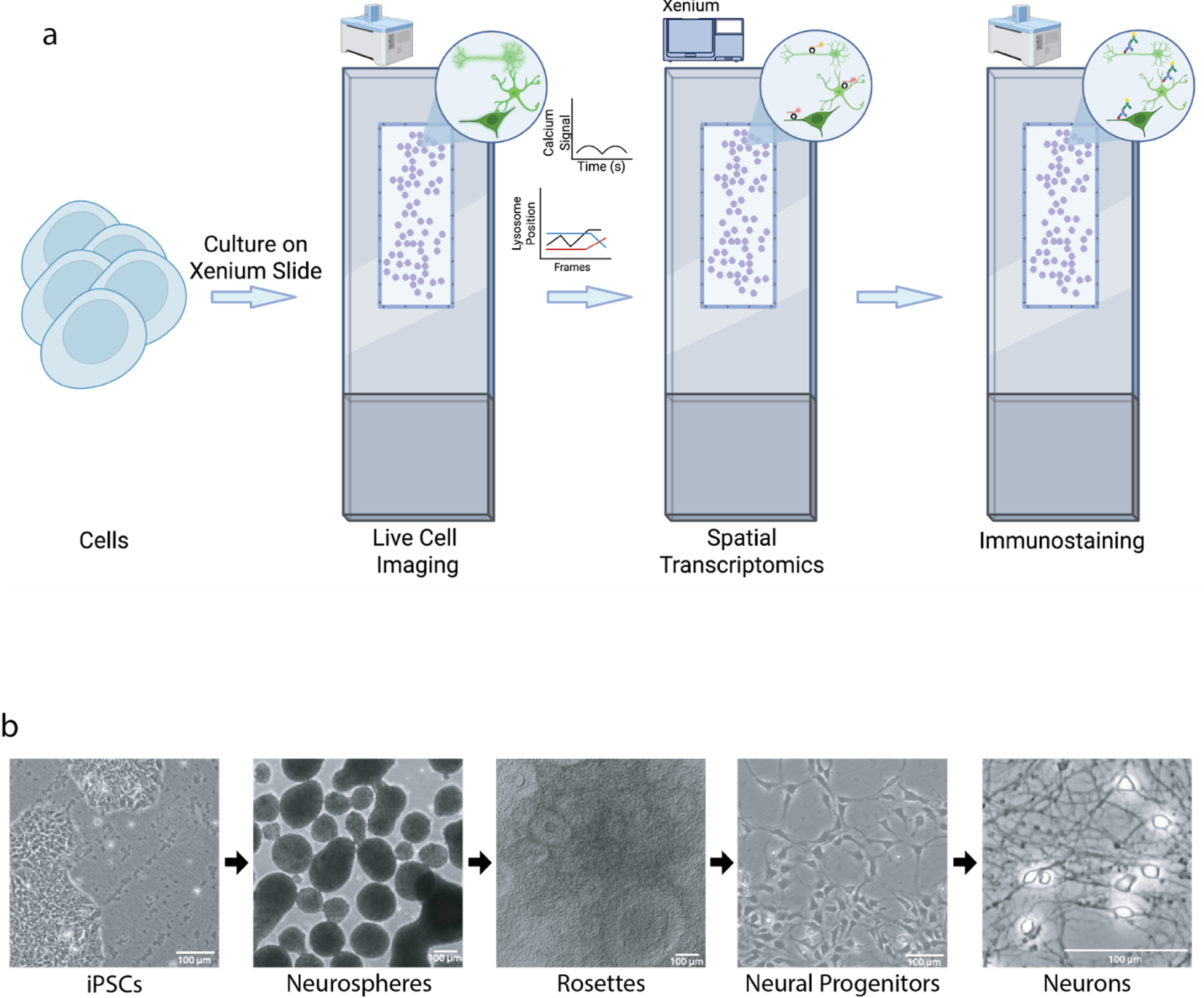
Diagram of VISTA-FISH workflow **a**. Diagram of VISTA-FISH approach. VISTA-FISH begins by culturing cells (such as neurons derived from iPSCs) on a glass slide compatible with spatial transcriptomics. A custom PDMS gasket placed on the slide ensures that cells grow only within the designated capture area. The VISTA-FISH workflow then performs three steps: (1) live-cell imaging using a confocal microscope to measure temporal phenotypes such as calcium transients indicating neuron activity; (2) spatial transcriptomic measurement of gene expression using the 10X Genomics Xenium Analyzer; and (3) immunofluorescent imaging. Image registration is then used to link the same cells in all three imaging rounds, providing a video, gene expression, and morphology for the same cells. **b**. Images of the stages in the dual SMAD inhibition protocol used to make neurons. Cells were plated onto the glass slide for VISTA-FISH at the neural progenitor stage (fourth image).

We chose to develop VISTA-FISH based on the 10X Genomics Xenium ST technology. Xenium offers a number of advantages: it is compatible with fixed cells; uses a glass slide that does not contain any cytotoxic materials; measures a relatively large panel of genes (up to 5,000); and allows the addition of custom padlock probes that can sensitively detect transcripts using junction-specific ligation and rolling-circle amplification. We faced two key challenges in developing VISTA-FISH: growing cells on the glass slide used for Xenium profiling, and registering live-cell images with ST data.

We first investigated how to culture cells on Xenium slides without inducing cytotoxicity or interfering with ST measurement. We found that polyethyleneimine (PEI) coating is an effective coating for neuron culture, allowing us to culture neurons for 4-6 weeks on the Xenium slide. An additional challenge is that each Xenium slide contains embedded fiducial marks, which are necessary for registering images acquired over multiple rounds of probe hybridization. If cells grow over the fiducials, the Xenium analyzer fails to properly register images, interfering with ST measurements. To prevent cells from growing over these fiducials, we designed a custom polydimethylsiloxane (PDMS) gasket that fits on the Xenium slide, with a cutout overlapping the spatial transcriptomic capture area (**Supplementary** Fig. 1a,b). The gasket is compatible with inverted microscopes, yet provides an effective barrier, ensuring cells grow only within the desired capture area and do not overgrow the fiducials. The gasket is removed after live-cell imaging and before loading the slide onto the Xenium analyzer, resulting in a rectangular lawn of cultured cells within the Xenium capture area.

We next sought to link live-cell imaging and Xenium ST data. This is possible in principle because cells remain in the same position during live-cell imaging and sequential probe hybridization on the Xenium analyzer. However, the live-cell imaging is performed using a different microscope than the Xenium Analyzer, and also with potentially different magnification. Thus, image registration is required to match the live-cell video from each cell with its gene expression. Although the fiducials on the Xenium slide could in principle be used to register images across microscopes, the Xenium Analyzer performs on-instrument preprocessing that automatically removes the fiducials from the images before exporting them. Thus, we sought an alternative registration approach. Instead of using the fiducials on the slide, we manually specified corresponding landmark points between the live-cell images and the spatial transcriptomic images. A single rigid transformation based on these landmarks was sufficient to accurately match the cell mask used for transcriptomic quantification with the live-cell images of the same cell (**Supplementary** Fig. 1c). Similarly, we registered immunostaining images acquired after spatial transcriptomics using manual landmarks.

### VISTA-FISH links calcium activity, gene expression, and morphology in differentiating neurons

As cells differentiate, sequential gene expression changes establish the molecular components that allow each cell to carry out specialized functions. For example, mature neurons express ion channels and neurotransmitter receptors that allow them to conduct action potentials and function in neural circuits by firing action potentials. However, because neuron activity is a temporal property that can only be measured in live cells, linking differences in neuron activity to molecular differences remains challenging. Patch-seq combines patch-clamp electrophysiology with scRNA-seq, but is technically difficult, labor-intensive, and low-throughput^7^. Conversely, live-cell imaging of calcium transients provides a high-throughput and quantitative means of measuring neuronal activity, but existing approaches for calcium imaging cannot simultaneously measure high-dimensional gene expression.

We used VISTA-FISH to investigate gene expression changes that occur as neurons develop into electrically active cells. As outlined in **Fig. 1b**, we differentiated human iPSCs via dual SMAD inhibition^8^, yielding a diverse population of cell types, including neural and glial progenitor cells, inhibitory and excitatory neurons, and astrocytes. For these studies, we used an iPSC line engineered to express a genetically encoded calcium indicator derived from green fluorescent protein (GCaMP)^9^. Differentiated cells at the neural progenitor stage were then plated onto 10X Xenium slides and cultured for 5 weeks. We imaged single-cell calcium transients at 3Hz using a spinning-disc confocal microscope (Yokogawa CQ1). After calcium imaging, we fixed and permeabilized cells and performed ST. To facilitate cellular segmentation, we added a nuclear dye (DAPI) in combination with immunocytochemical markers of intermediate filaments (vimentin) and cell surface proteins (cadherins) before Xenium imaging. We also immunostained for neurofilament (NFH) and microtubule-associated protein 2 (MAP2) to visualize axon and dendrite morphology, respectively. We then performed image registration to identify the same cells in the live-cell images, Xenium images, and immunofluorescence images.

Immunostaining of either MAP2 or VIM is shown, providing matched morphology information about the positions of dendrites or radial glia filaments for each cell. Colored dots represent individual transcript positions. Cell boundaries are shown as transparent masks.

Thus, for each cell, we obtained a video of GCaMP activity, images of several morphological markers, and the spatial positions of individual transcripts from 5,000 genes. We profiled a total of 155,237 cells, detecting a median of 803 transcripts/cell. After filtering cells with low transcript counts, we retained 81,547 cells for downstream analysis. Unsupervised clustering analysis identified 8 transcriptionally distinct groups of cells falling into two broad classes: neurons and non-neurons (**Fig. 2a-b, Supplementary** Fig. 3c). We observed a spectrum of cellular states expressing markers of excitatory and inhibitory neurons, astrocytes, neural progenitor cells, and radial glia (**Fig. 2b-d**).

**Figure 2.**
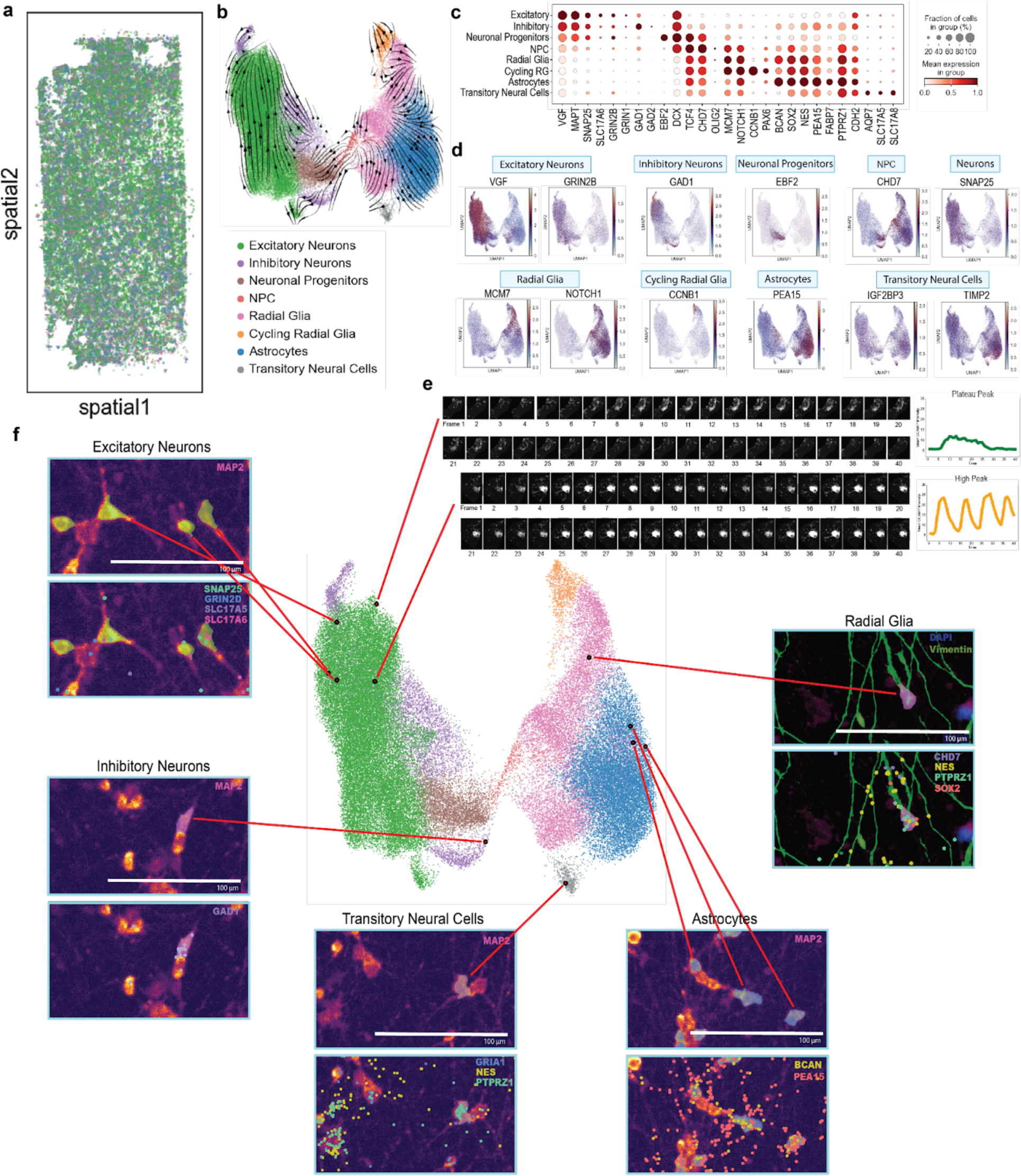
VISTA-FISH measures gene expression, morphology, and calcium activity in the same cells. **a**. Human iPSC-derived neurons (via dual SMAD inhibition) were cultured on a Xenium 10X Genomics glass slide. Each colored dot represents an annotated cell and the spatial coordinates of X (Spatial 1) and Y (Spatial 2). **b**. UMAP plot of single-cell gene expression data from VISTA-FISH and RNA velocity analysis. Each dot represents a cell. Arrows indicate differentiation trends. **c**. Dot plot of marker gene expression patterns from each cluster. **d**. Expression feature plots of marker genes from each cluster. **e**. Calcium signal from live imaging. Forty images were taken at 300 ms exposure. The graph represents the overall GCaMP intensity (y-axis) and the image frame number (x-axis). **f**. Paired morphology (top picture) and subcellular transcript localization (bottom picture) for selected cells.

To quantify the differentiation stages of the cells from their ST profiles, we performed RNA velocity analysis. Although RNA velocity analysis usually relies on measurements of spliced vs. unspliced transcripts to estimate transcription and splicing rates, nuclear vs. cytoplasmic localization from multiplexed FISH can instead be used to estimate nuclear export rates^10^. We therefore quantified nuclear and cytoplasmic localization of the transcripts from the 5,000 genes we measured, and used our previously published VeloVAE tool^11^ to estimate latent time and RNA velocity for each cell (**Fig. 2b, Supplementary** Fig. 2). The RNA velocity vectors and latent time are consistent with cell type annotations determined by ST profiling: RNA velocity analysis indicates that differentiation starts from radial glia-like cells and proceeds to either neuronal or non-neuronal cell states.

We further identified the top differentially expressed genes for each cluster and observed distinct gene expression patterns between neuronal and non-neuronal clusters (**Fig. 2c, d**). Neurons expressed relatively high levels of SNAP25, a presynaptic protein involved in neurotransmitter release. The majority of neuronal cells also displayed high expression of excitatory neuron markers (e.g., GRIN2B and VGF), as expected from dual SMAD inhibition. A subset of neurons also expressed GAD1, a marker of inhibitory neurons. In contrast, cells in the astrocyte cluster expressed glial markers such as PEA15, while cells in the radial glia cluster expressed MCM7 and NOTCH1. A small population of cells co-expressed cell cycle genes (e.g., CCNB1) and markers of radial glia. We also identified a small group of cells that expressed both glial and neuron markers, such as PTPRZ1, NES, SLC17A5, and SLC17A8. These cells also clustered near the astrocytes and displayed neurite-like processes (see below). Because of their overlapping markers and morphology, we refer to these cells as “transitory neural cells”. We also observed that some clusters were preferentially co-localized in space (**Supplementary** Fig. 2).

VISTA-FISH measures gene expression, live-cell images, and morphology in the same cells, providing an exciting opportunity to investigate the links among these cellular properties. Having characterized transcriptional differences among the cells from dual SMAD inhibition, we next investigated their morphology and activity. For each cell, we obtained both a transcriptomic cluster assignment and a video of time-dependent changes in GCaMP fluorescence, divided into 40 frames (**Fig. 2e**). In doing so, we observed significant cell-to-cell variation in calcium transients. For example, one excitatory neuron shown in **Fig. 2e** shows a single peak of GCaMP fluorescence during the recording period, while another shows multiple rapid spikes. We also observed dramatic differences in the morphology of cells across transcriptomic clusters (**Fig. 2f**). Cells in the excitatory neuron cluster possess a soma with characteristic pyramidal shape and display complex neurite morphologies as shown by MAP2 immunofluorescence. Cells in the inhibitory neuron cluster likewise show strong MAP2 positivity, but exhibit a longer, narrower soma than those in the excitatory neuron cluster. In contrast, cells in the radial glia cluster extrude vimentin (VIM)-positive filaments, while cells in the astrocyte cluster lack both VIM+ filaments and NFH/MAP2+ neurites. Transitory neural cells displayed both excitatory and astrocytic gene marker expression as well as VIM+ filaments.

### VISTA-FISH relates cell clusters based on calcium activity and gene expression

To further investigate the link between calcium activity and gene expression, we clustered cells by patterns of GCaMP fluorescence, then compared these patterns to gene expression cluster assignments for the same cells. Using the cell boundaries identified by the Xenium Explorer software, we extracted several features from the temporal GCaMP data, including median fluorescence intensity, number of peaks, area under the activity curve (AUC), peak amplitude, normalized peak amplitude (fold change above the median), and peak duration (**Supplementary** Fig. 3a-b). Grouping cells by these features resulted in six distinct clusters (**Fig. 3a-b**). Each cluster showed a distinctive pattern of activity, readily apparent from plots of fluorescence intensity over time (**Fig. 3a**). Most cells showed relatively invariant fluorescence levels (inactive, IA). Cells with more variable GCaMP transients fell into groups showing a single small peak (SP), multiple small peaks (MP), plateau peaks (PP), multiple moderate peaks (MoP), and multiple high peaks (HP). The distinctive properties of each cluster are also apparent from examining the distributions of features used for clustering (**Fig. 3b**). HP cells showed the most diverse features, including AUC above the mean, normalized peak amplitude, and peak amplitude.

**Figure 3.**
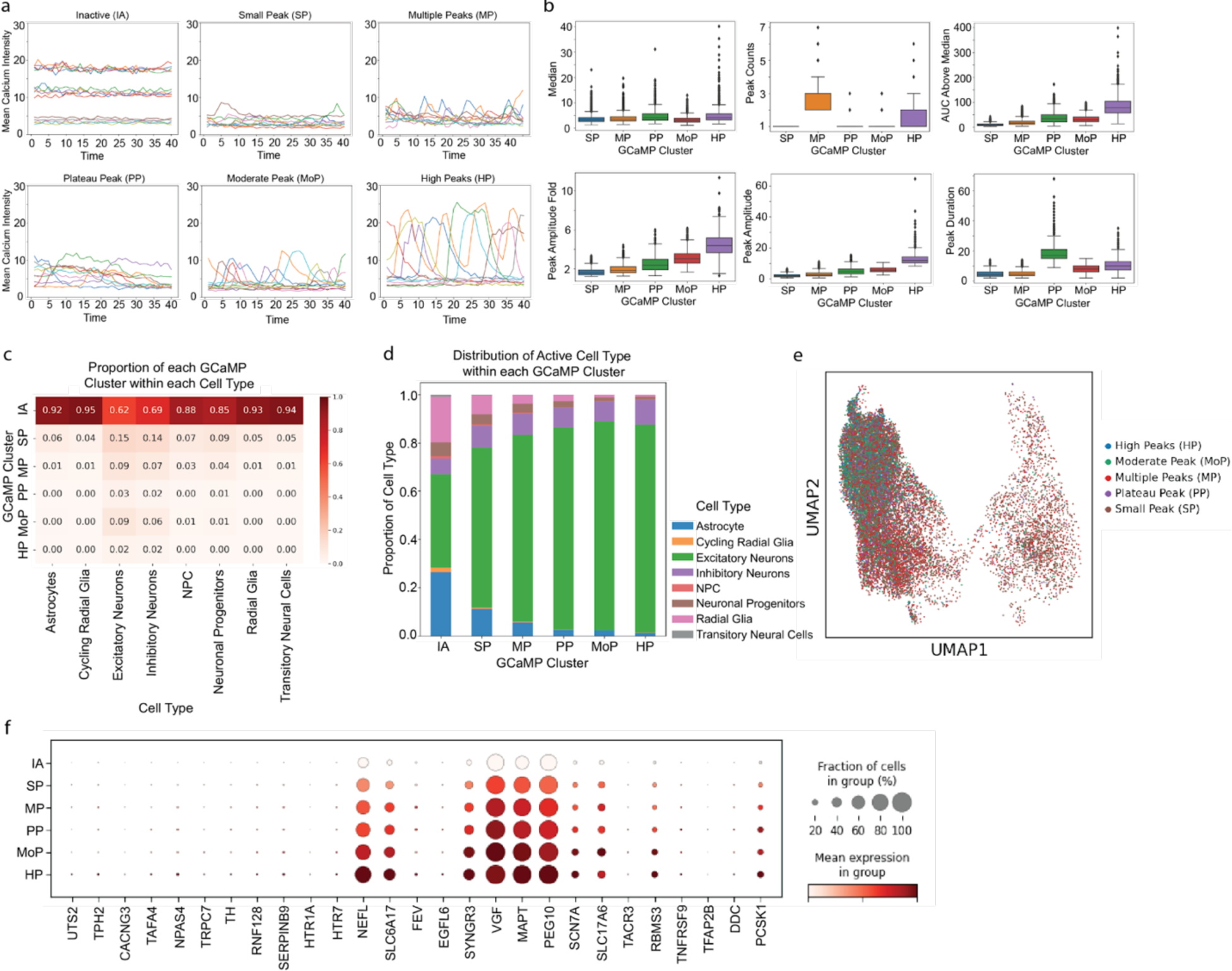
VISTA-FISH reveals how cells with distinct calcium activity distribute across transcriptional cell clusters. **a**. Calcium signal traces of cells grouped based on their pattern: Inactive (IA); small peak (SP); multiple peaks (MP); plateau peak (PP); moderate peak (MoP); high peaks (HP). Each line represents calcium signal from a cell recorded at 40 different timepoints (X-axis). **b**. Boxplots of summary statistics for GCaMP data across activity clusters. **c**. Proportions of activity clusters within each transcriptomic cluster. **d**. Proportions of transcriptomic clusters within each activity cluster. **e**. UMAP plot colored by activity clusters. Each dot is a single cell. The x and y coordinates are derived from transcriptomic data. The color indicates which activity cluster the cell belongs to. Cells from the IA cluster are not shown so that other colors can be distinguished. **f**. Dot plot of genes most differentially expressed across activity clusters.

From these analyses, we obtained two cluster assignments for each cell: a gene expression cluster and an activity cluster. We then examined the proportion of each activity cluster within each gene expression cluster (**Fig. 3c**). Most non-neuronal cells were assigned to the inactive cluster based on their GCaMP intensity (cycling radial glia: 95%; radial glia: 93%; astrocytes: 92%; NPCs: 88%). The few non-neuronal cells not in the inactive cluster were assigned to the SP and MP clusters, both of which exhibit low peak amplitudes. In contrast, 38% of cells in the excitatory neuron cluster and 31% of cells in the inhibitory neuron cluster belonged to the 5 activity clusters with more variable activity. We also quantified the proportion of each transcriptional cell cluster within each activity cluster. The cells in the clusters with variable activity almost exclusively belonged to the excitatory and inhibitory neuron clusters (**Fig. 3d**). A UMAP visualization of cells colored by their activity clusters confirms these relationships (**Fig. 3e**). We also asked whether the expression of specific genes predicts assignment to one or more activity clusters (**Fig. 3f**). The most significantly differentially expressed genes showed a gradient pattern, with lowest expression in the IA cluster, and increasing expression in clusters with more variable activity (SP, MP, PP, MoP, and HP clusters, in that order), implying that these genes regulate neuronal maturation and activity.

### VISTA-FISH enables prediction of calcium activity from single-cell gene expression

Our analyses indicated clear relationships between the transcriptional state of a cell and its activity. But which genes are most closely linked to variations in activity? To investigate this question, we used VISTA-FISH data to fit a linear regression model that predicts calcium activity from a sparse subset of genes. We chose the GCaMP intensity area under the curve (AUC) as the dependent variable for our regression analysis, because it effectively summarizes the total amount of activity measured during the period of observation. AUC is also a good choice for this analysis because it shows significant variation among our transcriptional cell clusters, particularly between neurons and non-neurons (**Fig. 4a-b**).

**Figure 4.**
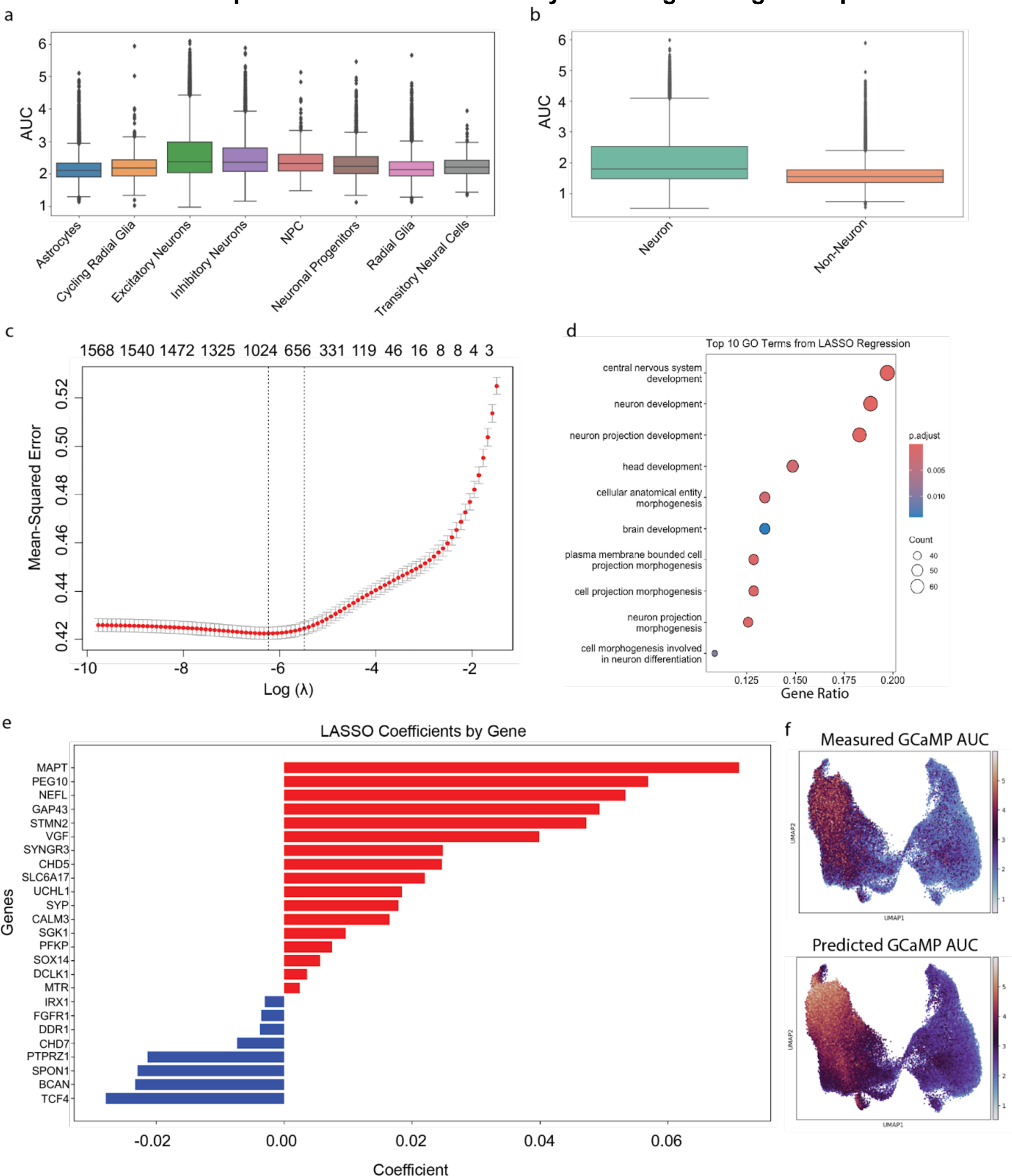
VISTA-FISH measures calcium signal patterns for each cell type and identifies neural activity-associated genes. **a**. A box plot of calcium signal using the area under the curve (AUC) for each cell type. **b**. Overall differences in AUC between neurons and non-neuronal cells. **c**. A plot of the Mean-Squared Error (MSE) and the number of genes associated with the calcium signal from RNA expression. The first and second dotted vertical lines represent minimum error and +1 standard error, respectively. **d**. Top-ranking Gene Ontology Biological Process terms associated with calcium signal. **e**. Top individual genes positively or negatively associated with calcium signal. **f**. UMAP of the measured (**top**) and predicted (**bottom**) AUC.

We employed linear regression with an elastic net penalty to predict single-cell GCaMP intensity AUC from gene expression. Through cross-validation, we selected an optimal model using 954 genes, which achieved a mean-squared error (MSE) of 0.42 with R^2^ = 0.25 (**Fig. 4c**). Notably, a much sparser model with only 26 genes attained a slightly higher MSE of 0.45 (**Fig. 4e**). Gene ontology analysis showed that these genes were enriched for factors involved with nervous system and neuron structural development, aligning with the observed heterogeneity in neural activity (**Fig. 4d**). Among the 26 genes, *MAPT* and *PEG10* emerged as the strongest positive predictors of neural activity, whereas *TCF4* and *BCAN* expression were inversely associated with GCaMP intensity (**Fig. 4e**). We then visualized the predicted vs. measured AUC from each cell on a UMAP plot (**Fig. 4f**). The predicted GCaMP signal showed a similar pattern to the measured intensity, with AUC increasing in the direction of neuronal differentiation, indicating that the model reasonably predicts activity from gene expression.

### VISTA-FISH detects neurite-enriched transcripts and differentiation-associated changes in nuclear localization

Subcellular transcript localization plays an important role in cellular function. For example, key neuronal proteins are translated from transcripts located in axons or dendrites, and RNA mislocalization is linked to neurodegenerative disease^12,13^. VISTA-FISH measures not only total gene expression levels per cell, but also the location of each transcript, providing an exciting opportunity to link RNA localization with cell morphology and temporal properties.

We first looked for transcripts enriched in neurites. Immunostaining for structural proteins, including MAP2 for dendrites and NFH for axons, enables the discrimination of these neuritic structures and facilitates a more precise characterization of neuronal morphology. We thresholded the MAP2 and NFH staining images to identify likely axon and dendrite regions, then masked out the soma locations identified from spatial transcriptomics. We then identified transcripts overlapping the remaining locations and performed a differential expression analysis to identify transcripts enriched in axons vs. soma and dendrites vs. soma (**Fig**. **5a, b**).

**Figure 5.**
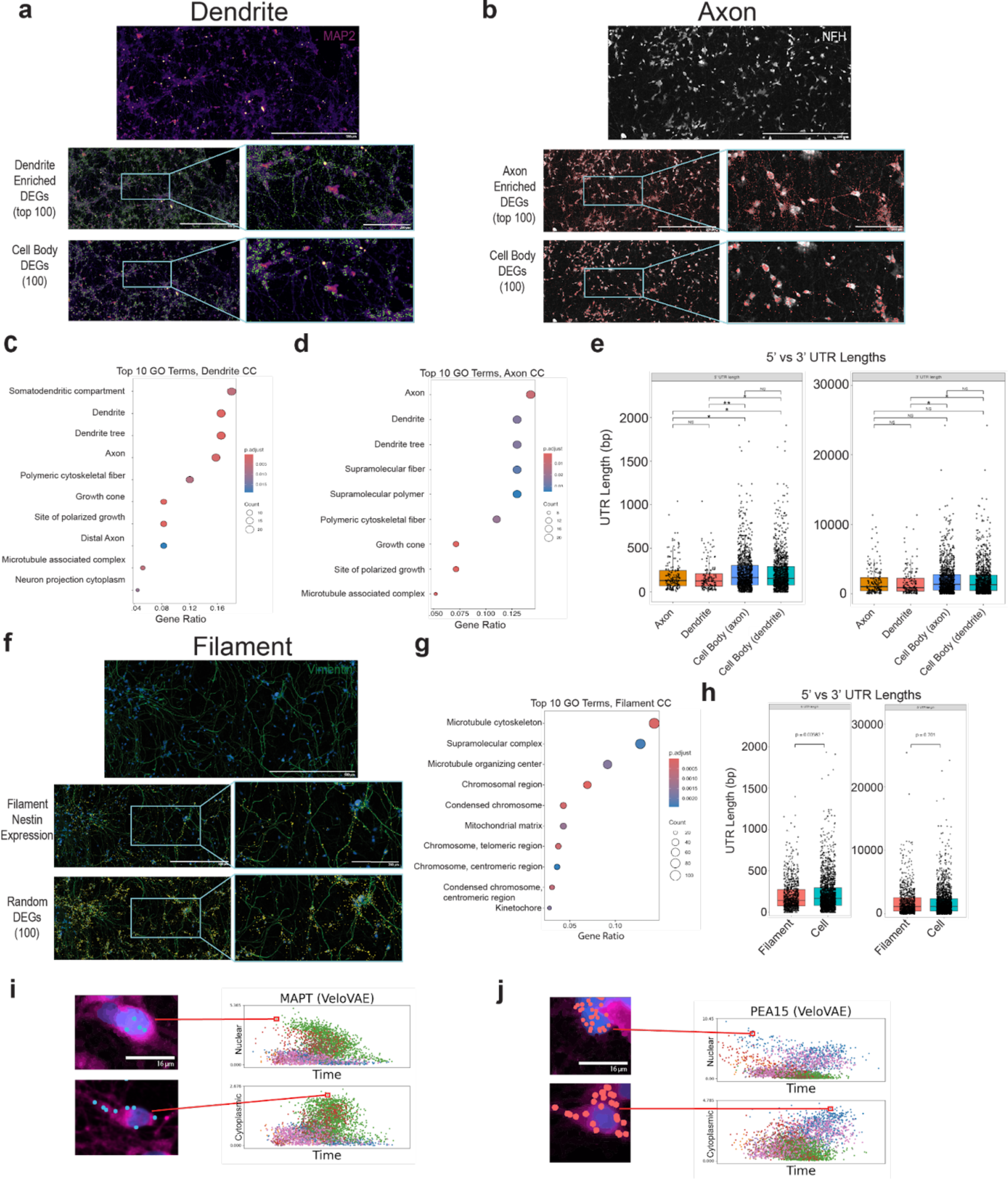
VISTA-FISH detects neurite-enriched transcripts and differentiation-associated changes in nuclear localization. **a.** Xenium immunofluorescence staining (IF) images of dendrites with MAP2 staining (purple), along with the top 100 dendrite-enriched and soma- enriched DEGs (green). The top panel represents the dendrites, while the bottom two show the top 100 dendrite-enriched DEGs and 100 soma-enriched DEGs, respectively. **b.** Images of neurons with axons stained with NFH (white), and top 100 axon-enriched DEGs panels (red), followed by top 100 soma-enriched DEGs. **c,d.** Dot plot of the top 10 gene ontology cellular component terms in dendrites **(c)** and axons **(d)**. The color indicates the enrichment P-value, and the diameter of the dots indicates the number of genes overlapping the GO term. **e.** Boxplot of 5’ (left) and 3’ (right) UTR lengths of genes enriched in axons, dendrites, soma, and cell bodies. **f.** Xenium slide images of Vimentin+ (green) radial glial filaments. Yellow dots represent the location of the Nestin transcript (middle panel) and soma-enriched genes (bottom panel). **g.** Dot plot of the top 10 cellular component terms enriched in genes with filament-enriched transcripts. **h.** Boxplot of 5’ and 3’ UTR lengths in filament-enriched and soma-enriched transcripts. **i,j.** Xenium IF images of neurons with transcript locations of MAPT (left panel) and **(j)** PEA15 (right panel) inside the nucleus (blue) and cytoplasm (pink). Scatter plot summarizing RNA velocity analysis, which indicates a change in nuclear vs. cytoplasmic localization (y-axis) over latent time (x-axis).

We detected 138 genes with dendrite-enriched transcripts and 159 genes with axon-enriched transcripts. Plotting the locations of individual transcripts from the top 100 dendrite-enriched and axon-enriched genes confirmed that these transcripts are indeed preferentially localized within neurites (**Fig**. **5a, b**). Gene ontology analysis indicated that many dendrite- and axon-enriched transcripts encode proteins that are themselves located within these subcellular compartments (**Fig. 5c, d**). For example, the top cellular component (CC) term for dendrite-enriched genes is “dendrite” and the top CC term for axon-enriched genes is “axon”. This result was not necessarily expected a priori due to the distinction between transcript localization (detected by our analysis) and protein localization (annotated in the GO CC terms). One potential explanation is that many proteins in dendrites and axons are locally translated from RNA in the same compartment, rather than undergoing transport to the axon or dendrite following translation within the soma^14^. We also observed moderate overlap between axon-enriched and dendrite-enriched genes, possibly due to shared molecular requirements for each compartment.

We also compared the lengths of the annotated untranslated regions (UTRs) among dendrite-enriched, axon-enriched, and soma-enriched transcripts. The transcripts abundant in dendritic and axonal compartments possess significantly shorter 5’ UTRs than those predominantly localized to the soma, and dendrite-enriched transcripts also show significantly shorter 3’ UTRs (**Fig**. **5e**). This finding is consistent with prior evidence suggesting that transcripts with shorter UTRs exhibit enhanced stability, which may facilitate efficient transport along the axon^15–17^.

We similarly identified transcripts enriched in vimentin (VIM) + processes of radial glia-like cells (**Fig**. **5f**). A total of 738 genes showed significant enrichment in VIM+ processes compared to the soma. *Nestin* transcripts showed particularly striking enrichment in VIM+ processes, consistent with previous data demonstrating local translation of Nestin within radial glia and astrocyte VIM+ filaments^18,19^. Moreover, filament-enriched transcripts tend to encode genes with microtubule-associated CC terms (**Fig**. **5g**). The 5’ UTRs of filament-enriched transcripts are slightly shorter than soma-enriched genes, though this effect is less pronounced than in neurite-enriched transcripts (**Fig**. **5h**). In contrast, the 3’ UTRs of filament-enriched transcripts showed similar lengths to the UTRs of soma-enriched transcripts.

We further investigated changes in RNA localization linked to cell differentiation. Our RNA velocity analysis based on nuclear vs. cytoplasmic localization predicted cell differentiation stages concordant with our transcriptomic cluster assignments (**Fig. 2b**). This suggests that many of the 5,000 genes we measured may undergo coordinated changes in nuclear export during differentiation. Indeed, we identified multiple genes with clear shifts in nuclear localization as they differentiated toward either a neuronal or astrocytic fate. For example, *MAPT* transcripts demonstrated predominant nuclear localization in immature excitatory neurons, while the same transcript was largely cytoplasmic in mature excitatory neurons (**Fig**. **5i**). This is consistent with rapid *MAPT* transcription in immature neurons that precedes nuclear export and translation in more mature cells. Similarly, *PEA15* transcripts were concentrated in the nucleus of immature astrocytes, but exhibited cytoplasmic accumulation in more differentiated astrocytes (**Fig. 5j**).

This suggests that *PEA15* transcripts appear in the nucleus first as transcription begins early in astrocyte differentiation, then transcripts appear also in the cytoplasm as nuclear export occurs and transcription continues as the cells differentiate.

### VISTA-FISH reveals changes in lysosome movement, cell morphology, and gene expression after genetic perturbation

Targeted genetic perturbation, such as CRISPR genome editing or CRISPR interference (CRISPRi)^20–23^, can be a powerful tool for probing the molecular mechanisms that underlie cellular phenotypes. Even so, investigating phenotypes with a temporal component remains challenging with CRISPR screening approaches. For example, lysosomal and mitochondrial transport are crucial for proper neuronal function^24–27^. However, a pooled CRISPR screen for this sort of temporal phenotype is challenging with existing approaches, because one must choose between either live-cell assays that can measure organelle movement, and static but high-dimensional molecular measurements of gene expression. Similarly, optical pooled screening can link live-cell images of a cell with genetic perturbations, but has not been used to simultaneously measure a live-cell phenotype and how the perturbation alters gene expression^28,29^. An ideal screening approach would be capable of measuring how lysosome movement and gene expression within the same cell change after genetic perturbation.

Thus, we extended VISTA-FISH to enable pooled CRISPR screening by developing a strategy to detect single-guide RNAs (sgRNAs) using custom Xenium probes. Briefly, we designed custom padlock probes with two 25-bp arms (**Fig. 6a**). Both probe arms must bind and be ligated to enable rolling circle amplification and target transcript identification. To detect CRISPR-Cas9 sgRNAs, we designed one arm of the probe to target the protospacer sequence unique to each sgRNA, and the other arm to target an adjacent region that is shared across all sgRNAs. Crucially, this approach allows us to detect sgRNAs without modifying the sgRNA vector or performing *in vitro* transcription—key advantages compared to recent papers that detected sgRNAs with FISH^30^. Additionally, using the Xenium chemistry, we can profile 5,000 genes in addition to the sgRNAs.

**Figure 6.**
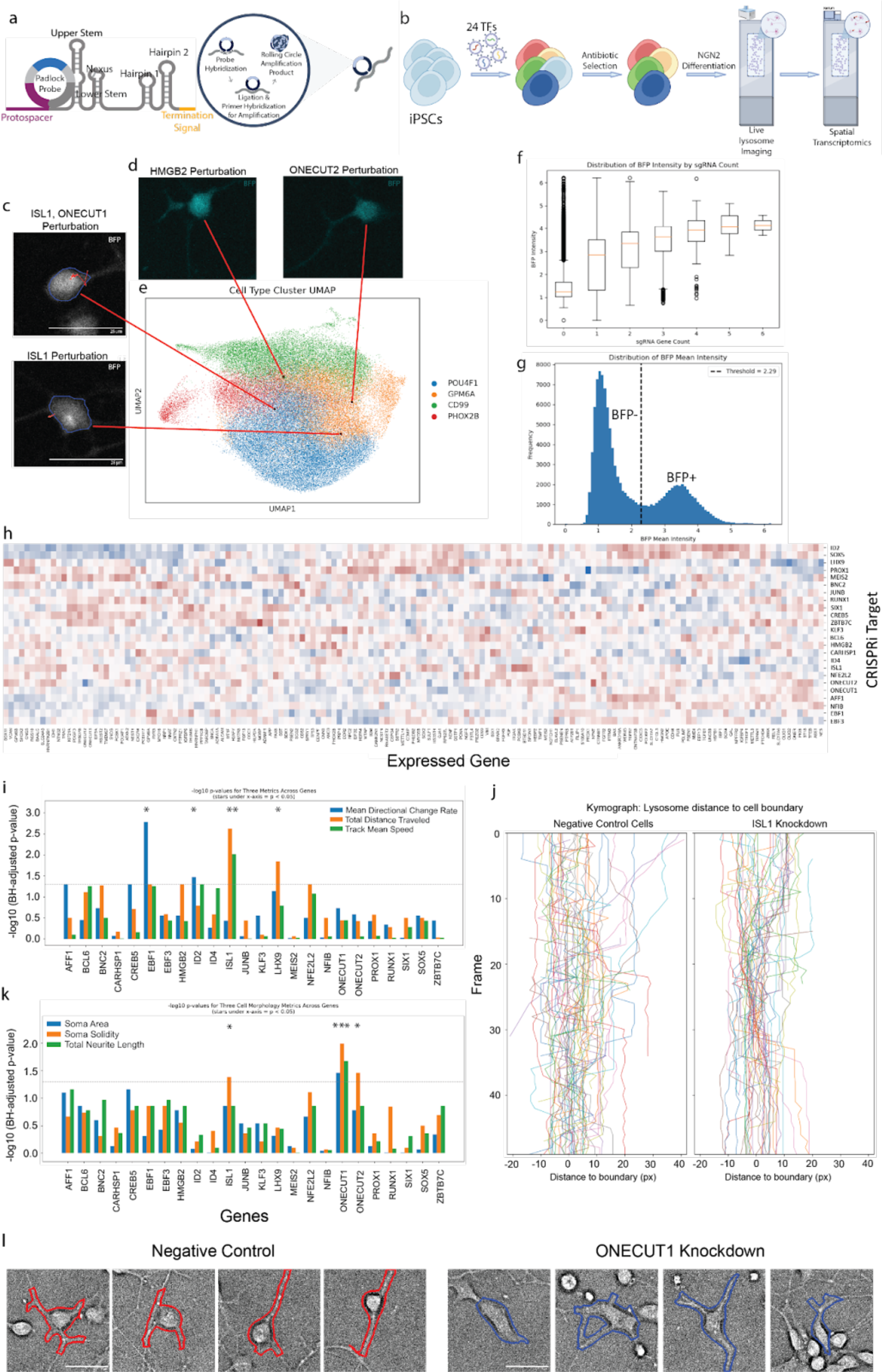
VISTA-FISH detects changes in lysosome dynamics and cell morphology in a pooled CRISPR interference screen. **a**. Diagram of VISTA-FISH approach for detecting CRISPR sgRNAs. One arm of each padlock probe targets the unique protospacer sequence of each sgRNA, while the other arm targets the adjacent shared sgRNA sequence. Probe ligation and rolling circle amplification allow sensitive and specific sgRNA detection without requiring additional steps such as *in vitro* transcription. **b**. Schematic of pooled CRISPRi screening experiment in human iNeurons. **c-e**. VISTA-FISH measures brightfield images of lysosome movement. Red lines indicate the paths taken by individual lysosomes. **d**. Images of two neurons with CRISPRi gRNA expressing blue fluorescent protein. **e.** UMAP of the 59,466 cells measured in the experiment. Red lines indicate the UMAP coordinates of the cells. **f**. A box plot of the distribution of BFP intensity by sgRNA count. **g.** Histogram of log(BFP) for cells profiled by VISTA-FISH. The vertical dotted line indicates the threshold used to separate BFP+ and BFP- cells. **h.** Heatmap of 148 genes (columns) showing significant differential expression across the 24 CRISPRi perturbations (rows). **i.** Bar plot of the -log10 transformed and Benjamini-Hochberg adjusted p-value of lysosomal directional change rate, distance traveled, and mean speed in each CRISPRi target compared to negative control. \**FDR*<0.05. **j.** Kymograph of lysosome distance to cell boundary for 50 randomly selected lysosome tracks in both the negative control and ISL1 knockdown groups. Lysosomes were selected from the 48th-60th percentiles of total distance traveled in each group to ensure representative sampling. **k.** Bar plot of the -log10 transformed and Benjamini-Hochberg adjusted p-value of soma area, solidity, and total neurite length in each CRISPRi target compared to negative control. \**FDR*<0.05. **l.** Representative images of soma and neurite boundary outlines in cells with negative control (left) or ONECUT1 (right) sgRNAs overlaid on brightfield images.

We then designed a VISTA-FISH experiment to profile the effects of gene knockdown on lysosome trafficking and neuronal morphology (**Fig. 6b**). We chose to perform the screen in human iNeurons, a model of glutamatergic neurons differentiated by induced expression of the master transcription factor NGN2 in iPSCs. To enable CRISPRi, we used iPSCs that also stably express dCas9 fused to the transcriptional repressor KRAB. Based on previous transcriptomic data^31^, we selected 24 transcription factors (TFs) highly expressed in iNeurons. We used a lentiviral vector to introduce a pooled sgRNA library targeting all 24 TFs (5 sgRNAs per gene) (**Fig. 6b**). The sgRNA vector also encodes a blue fluorescent protein (BFP) reporter and a puromycin resistance cassette, allowing both positive and negative selection of transduced cells. Following antibiotic selection, cells were differentiated into iNeurons, then plated onto the Xenium slide. We then added a lysosome dye (lysotracker-red) and performed live-cell imaging of lysosomes and BFP fluorescence, enabling direct identification of sgRNA-positive and - negative cells through protein expression. After imaging, we immediately fixed and permeabilized cells, then proceeded to Xenium spatial transcriptomics.

We thus obtained a VISTA-FISH dataset with live-cell imaging of lysosomes and BFP cells for 50 frames, in addition to single-cell gene expression and sgRNA identity in fixed cells (**Fig. 6c- d**). We obtained high-quality gene expression data, detecting a total of 59,466 cells with a median of 1033 transcripts per cell. To quantify lysosome movement from live-cell images of lysotracker dye, we identified and tracked lysosomes using TrackMate^32^, obtaining a trajectory for each lysosome. We then registered the lysosome tracks to Xenium cell masks, allowing us to link the gene expression profile of each cell with lysosome tracks that overlapped the soma during all or part of the live-cell imaging period (**Fig. 6c**). Live-cell BFP fluorescence allowed us to validate our probe-based detection of sgRNAs (**Fig. 6d**). After filtering to remove cells with low gene expression counts, 59,466 cells were used for unsupervised clustering and UMAP visualization, revealing four distinct transcriptomic clusters (**Fig. 6e**), consistent with previously reported transcriptional subtypes of iNeurons^31^. We then compared sgRNA abundance within each cell to BFP intensity measured by confocal microscopy, which showed strong concordance: cells with the highest BFP signal harbored multiple distinct sgRNAs, while cells with low BFP intensity showed fewer sgRNAs (**Fig. 6f**). By classifying cells as BFP+ or BFP- and comparing these labels to the VISTA-FISH sgRNA counts, we found that our probes detected sgRNAs with 62.2% sensitivity and 85.7% specificity (**Fig. 6g**).

We next investigated how gene expression, lysosome movement, and morphology differed across the 24 genetic perturbations included in the screen. We observed statistically significant differential expression (FDR < 0.05) across perturbation conditions for 148 out of 358 genes profiled (**Fig. 6h**). To evaluate functional consequences of TF knockdown, we quantified lysosome dynamics using multiple metrics calculated from each lysosome track, including confinement ratio, mean directional change rate, linearity of forward progression, total distance traveled, track displacement, and track mean speed (**Supplementary** Fig. 4b). Out of the 24 TFs we screened, knockdown (KD) of 4 TFs resulted in statistically significant (FDR < 0.05) reductions in lysosome motility (**Fig. 6i, Supplementary** Fig. 4a). *ID2* and *EBF1* KD resulted in reduced lysosome directional change rate compared to cells transduced with negative control sgRNAs. Additionally, *ISL1* KD led to decreases in lysosome speed and total distance traveled, while *LHX9* KD similarly reduced the total distance traveled. We further visualized the lysosome tracking data using kymographs, confirming differences in lysosome movement between cells transduced with targeting vs. non-targeting sgRNA (**Fig. 6i** and **Supplementary** Fig. 4c). These effects all imply disrupted lysosomal trafficking, potentially impairing local degradation processes critical for neuronal maintenance^33,34^. Consistent with our results, both *LHX9* and *ISL1* are essential regulators of neuronal differentiation, neurogenesis, and axon guidance and growth^35–38^.

Finally, we used CellPose-SAM to segment soma and neurites, allowing us to compare iNeuron cell morphology across perturbations^39^. To facilitate accurate segmentation, we fine-tuned two separate models to identify neurites and soma, then merged neurite and soma masks to link neurites with soma that they touch or overlap (**Supplementary** Figs. 5-8). We then used CellProfiler to calculate a variety of morphometric features quantifying cell shape and neurite complexity, including soma area and shape, and neurite area and shape^40^. By comparing these features across our perturbations, we identified three perturbations with significant effects on cell morphology: *ONECUT1*, *ONECUT2*, and *ISL1* (**Fig. 6k, Supplementary** Fig. 9a).

*ONECUT1* KD resulted in significant changes to soma area, soma shape, and total neurite length (**Fig. 6k,l, Supplementary** Fig. 9d-f). Cells with *ONECUT2* and *ISL1* KD also showed significantly different soma shape (**Supplementary** Fig. 9b,c). Our findings are consistent with a previous study that found overexpression of *ONECUT1* and *ONECUT2* in human fibroblasts induced a neuron-like morphology^41^.

## Discussion

In this study, we developed VISTA-FISH, a platform that combines live-cell imaging with subcellular-resolution measurement of gene expression and detection of CRISPR guide RNAs. We used VISTA-FISH to measure gene expression, morphology, and calcium transients–a proxy for neuronal activity–in the same cell. We also performed pooled CRISPRi screening to perturb gene function and mapped resulting changes in lysosome activity and morphology.

VISTA-FISH combines readily available commercial platforms for microscopy and spatial transcriptomics, which may make it significantly easier for individual labs to adopt than approaches requiring custom sequential FISH setups^28,42^. A recently published approach, Perturb-Fish, combined imaging spatial transcriptomics with CRISPRi perturbation, paired with a functional readout using multiplexed error-robust fluorescence in situ hybridization (MERFISH)^30^. Perturb-Fish requires modifying the CRISPRi sgRNA vector and performing in vitro transcription using T7 polymerase. In contrast, our strategy detects sgRNAs without any required modifications to the sgRNA vector or additional protocol steps. Additionally, we profiled 5,000 genes, whereas the Perturb-Fish platform was limited to ∼500 genes.

Our study leaves several areas of improvement in future work. First, we measured a pre-designed panel of 5,000 genes. The panel was designed for broad applicability across tissue types, rather than tailored for brain-related genes. More tailored gene panels may identify more phenotype-relevant molecular mechanisms. Second, we performed live-cell imaging for a limited amount of time. For example, in our analysis of neuron activity, approximately 70% of cells identified as excitatory neurons based on gene expression did not exhibit detectable peaks within our imaging period–likely due in part to the relatively short period of observation. This is not an inherent limitation of the technology, and temporal phenotypes requiring longer imaging times will be interesting to explore in future studies. Finally, VISTA-FISH relies on custom probes to detect CRISPR sgRNAs. In the current study, we relied on 10X Genomics to synthesize these probes for us, which limited us to 125 targets. This is not an inherent limitation of the technology, however.

VISTA-FISH is a broadly useful tool that promises to enable a broad range of studies in molecular and cellular biology. By bridging functional phenotyping with transcriptomic profiling, VISTA-FISH overcomes a major limitation in current single-cell approaches. VISTA-FISH can be readily applied to study additional temporal phenotypes, including mitochondrial dynamics, cell migration, protein metabolism, and signal transduction. Finally, we hope that VISTA-FISH can help elucidate how genetic and environmental factors contribute to temporal cell phenotypes underlying complex diseases, such as neurodegeneration and psychiatric disorders.

## Methods

### iPSC maintenance

iPSCs stably expressing GCaMP6f were cultured in mTeSR1 (STEMCELL Technologies, 85850) on dishes with Matrigel (Corning, 354230). When iPSCs reached 70-80% confluency, cells were passaged by washing with DPBS (Gibco, 14190144), and dissociated with Accutase (Gibco, A1110501) at 37°C for 5 minutes. Accutase was then diluted 1:2 in DPBS, and the cells were collected in conical tubes and centrifuged at 300xg for 3 minutes. The supernatant was aspirated and resuspended with mTeSR1 supplemented with ROCK inhibitor (Stemcell, NC1678100). The cells were counted and plated onto Matrigel-coated plates at the appropriate number.

### Culturing Cells on the Xenium Slide

The Xenium slide 10X (Genomics) and cassette were submerged and sterilized in 70% ethanol for 15 minutes. Ethanol was removed from the dish, and the slide and cassette were air-dried for 20 minutes in the biosafety cabinet. The Xenium slide and cassette were treated with UV light for 20 minutes. We designed custom polydimethylsiloxane gaskets (fabricated by GRACE BIO-LABS) to cover fiducials on the Xenium slide (Figure S1). The gasket was placed on the slide and assembled in the Xenium cassette. 0.1% polyethyleneimine (PEI, Sigma, 306185) was added to coat the Xenium slide. The lid was placed on the cassette and incubated overnight at 37°C in a cell culture dish. The coating mixture was aspirated the next morning, rinsed three times with DPBS, and air-dried for 20 minutes. For plating NPCs from the dual SMAD protocol, the slide was coated with 20 μg/mL laminin (Sigma, L2020) for 2 hours at 37°C.

### Neuronal differentiation from iPSCs via dual SMAD inhibition

gCaMP6f-iPSCs were cultured on Matrigel-coated plates in mTeSR1 for 4 days. Cell clusters were lifted using L7 Passaging Solution (Lonza, FP-5013) and suspended in embryoid body (EB) medium ((DMEM/F12 (Gibco, 11320033), 20% KnockOut Serum Replacement (Gibco, 10828028) 1% NEAA (Gibco, 11140050) 2 mM GlutaMAX (Gibco, 35050061), 0.1 mM B-mercaptoethanol (Sigma, M6250)) supplemented with 10 uM ROCK inhibitor Y-27632 (Tocris, 1254, first day only), 2 uM dorsomorphin (Cayman Chemical, 866405-64-3) and 2 uM A-83 (STEMCELL Technologies, 72022) in low adherence plates for 4 days. On day 5, the medium was replaced with neural induction medium (NIM) consisting of DMEM/F12, 1X N2 supplement (Gibco, 17502048), 1% NEAA, 2 mM GlutaMAX, and 2 μM cyclopamine (Cayman Chemical, 4449-51-8) for another 3 days. Neurospheres were plated onto culture dishes coated with 100 μg/mL poly-L-ornithine (PLO, Sigma, P4957) and 10 μg/mL laminin in NIM. The centers of rosettes were cut and manually lifted after 7 days and replated onto PLO/laminin-coated dishes. After another 7 days, rosettes were manually dislodged and dissociated with Accutase (Corning, 25-058-CI) into single NPCs, which were seeded onto PLO/laminin-coated plates in NPC maintenance medium (Neurobasal Medium (Gibco, 21103049)), 1X B27 without Vitamin A supplement (Gibco, 12587010), 2 mM GlutaMAX, 1% NEAA, 20 ng/ml bFGF (PeproTech, 100- 18B), and 2 uM cyclopamine (Cayman Chemical, 11321).

NPCs were dissociated into single cells with Accutase, washed with DPBS, resuspended in NPC maintenance medium without bFGF, and plated onto PEI- and laminin-coated Xenium slides. The next day, an equal volume of complete BrainPhys (BrainPhys basal medium (STEMCELL Technologies, 05790), 1X SM1 supplement, 1X N2, 10 uM bucladesine (Cayman Chemical, 16980-89-5), 20 ng/mL BDNF (PeproTech, 450-02), and 20 ng/mL GDNF (Preprotech, 450-10)) containing 200 nM compound E (STEMCELL Technologies, 73952) was added. After three days, a half-volume media change was performed with complete BrainPhys. From this point, cells were grown for 5 weeks with twice-weekly half-volume media change with complete BrainPhys.

### iPSC-derived iNeuron differentiation

iPSCs-derived neurons (iNeurons) were differentiated from WTC11-iNgn2-dCas9 (generously gifted by Dr. Yin Shen from UCSF) using the published protocol^43^. On Day 0 of iNeuron differentiation, induction medium (IM, DMEM/F12, 1x of N2 supplement, 1x of non-essential amino acids (Gibco, 11140050), 1x of GlutaMAX) was prepared with doxycycline (2 mg/mL, 1000x) and ROCK inhibitor (10 mM, 1000x). Briefly, iPSCs were dissociated using Accutase, centrifuged at 300g for 3 min at room temperature, aspirated the supernatant, and resuspended the cells in IM supplemented with ROCK inhibitor and doxycycline. An appropriate number of cells were plated onto Matrigel-coated plates and incubated overnight at 37°C. On Days 1 - 3, the old medium was aspirated and washed with DPBS. Fresh induction medium supplemented with doxycycline was added to the plates.

On day 3 of iPSC-derived iNeuron differentiation, the cells were washed with DPBS and dissociated using Accutase for 5 minutes. Accutase was diluted with 1:2 DPBS and centrifuged for 3 minutes at 300g. Cells were resuspended in cortical medium (BrainPhys (STEMCELL Technologies, 05790), 1 μg/mL of Laminin, 10 ng/mL of BDNF, 10 ng/mL of NT-3 (PeproTech, 450-03), and 1x of B27 supplement (Gibco, 17504044)). Cells were seeded at 1 x 10^5^ in 250 μL of cortical medium supplemented with 1% pen/strep antibiotic on the sample area and incubated at 37°C, 5% CO2. An additional 750 μL of cortical medium was added the next day. The media was changed every other day.

### Molecular cloning and gRNA library

Lentiguide(10x)-BFP-Puro was generously gifted by Dr. Gary Hon from UT Southwestern Medical Center. We chose to perturb 24 TFs showing high expression in a previously published iNeuron single-cell RNA-seq dataset^31^. We selected 5 previously validated guide RNAs^44^ per target gene, plus 5 additional non-targeting guides. The oligonucleotide pool (Twist Bioscience) was PCR amplified and cloned using Gibson assembly into the single gRNA expression plasmid. The transformed plasmids were electroporated into Endura DUOs electrocompetent cells (LGC Biosearch Technologies, 60242-1). Transformed cells were added to LB broth supplemented with Ampicillin and incubated for 12-16 hours at 37°C in a shaker, and then proceeded to Maxi prep for plasmid DNA extraction (QIAGEN, 12162). Whole plasmid sequencing (Plasmidsaurus) was performed to verify that gRNAs were cloned successfully.

### HEK293T culture and lentivirus production

HEK293T cells were cultured in DMEM (Gibco, 11995065) supplemented with 10% FBS. Once the cells reached 80% confluency, they were dissociated using TrypLE Express (Gibco, 12604013) for 3 minutes at 37°C. TrypLE was diluted with DPBS and centrifuged for 3 minutes at 300g. Cells were counted and seeded onto a plate at the appropriate number.

For lentivirus production, 10-cm dishes were seeded with 7 x 10^6^ HEK293T cells in 12 mL of lentivirus packaging medium (Opti-MEM I (Gibco, 31985070), 5% FBS, 200 μM sodium pyruvate (Gibco, 11360070)). The next day, the transfection was performed using the following protocol by combining transfection mixes A and B. Transfection mix A contained 10 μg CRISPRi library and 26 μL of Lentiviral Packaging Mix (MISSION, SHP001) was mixed with 1.5 ml of Opti-MEM I and 36 ul of P3000 enhancer reagent (Invitrogen, L3000015). Transfection mix B contained 1.5 ml of Opti-MEM I and 42 μL of Lipofectamine 3000 reagent (Invitrogen, L3000015). Transfection mixes A and B were combined and incubated for 20 minutes at room temperature. During incubation, 6 mL of lentivirus packaging medium was removed from the 10 cm plate. The transfection mixes A and B were gently added and mixed. Six hours later, the Lipofectamine-containing medium was carefully removed and replaced with fresh lentivirus packaging medium. Lentivirus-containing medium was collected after 24 hours and stored at 4 °C. 12 ml of fresh lentivirus packaging medium was carefully added. Lentivirus-containing medium was collected again the next day. Combined lentivirus-containing medium (approximately 24 ml) was filtered through a 0.45 μm filter to remove cell debris into a new 50 ml conical tube. After filtering, ⅓ volume of Lenti-X Concentrator (Takara, 631231) was added to the filtered solution, mixed well, and stored at 4°C overnight. Following incubation, the solution was centrifuged at 4°C for 45 minutes at 1,500g. The supernatant was removed carefully, and the virus pellet was resuspended with 1 mL mTeSR and stored at -80°C until further analysis.

### GCaMP live-cell imaging

Using the manufacturer’s protocol, the media was switched to BrainPhys Imaging Optimized Medium (STEMCELL Technologies, 05796) the day before the live imaging. Briefly, ¾ of the medium was removed, and the same volume of fresh BrainPhys Imaging Optimized Medium with supplement was added. After 30 minutes of incubation at 37°C, ¾ of the medium was removed and the same volume of fresh medium mixture was added. After overnight incubation at 37°C, half of the medium was removed, and the same volume of the fresh medium mixture was added on the day of live cell imaging.

Calcium imaging was acquired using the CellVoyager CQ1 (YOKOGAWA). GFP images (400- ms exposure) were acquired over time-lapse, one Z-plane with 40 frames for each field of view (FOV) with laser enhancement at 10x. Acquired images were stitched together using Fiji. The cells were kept in PBS-T (0.05% Tween-20) at 4°C until further analysis.

### Lysosome Imaging

iNeurons were incubated for 30 minutes with 50 nM LysoTracker Red DND-99 (Invitrogen, L7528) in cortical medium at 37°C, the day of live imaging. The LysoTracker containing the cortical medium was replaced with the fresh cortical medium. Red fluorescent images were acquired over time-lapse using 300-ms exposure and 50 frames for each FOV at 20x for lysosome imaging using the CellVoyager CQ1. Images were stitched together using a Python script. The cells were kept in PBS-T (0.05% Tween-20) at 4°C until further analysis.

### Xenium Spatial Transcriptomics

The Xenium cell culture protocol was used. Briefly, cells on the xenium slide were washed with 1X PBS after live imaging. 1X PBS was removed carefully, and 4% paraformaldehyde (PFA) was added to fix cells and incubated at room temperature for 30 minutes. PFA was removed, and cells were washed three times with 1X PBS. PBS was removed, and pre-chilled 100% methanol was added and incubated at -20°C for 1 hour. Cells were washed with 1X PBS three times and hydrated with 1X PBS. Room temperature Probe Hybridization Mix was added to cells and incubated overnight. Cells were washed twice with PBS-T, and the Post-Hybridization Wash Buffer was added. The cells were washed with PBS-T three times. The Ligation Mix was added to cells and placed on the thermal cycler for two hours at 42°C. After ligation, the cells were washed three times with PBS-T. Amplification was performed by adding the Amplification Enhancer Master Mix and was placed on the thermal cycler for 2 hours at 4°C. The cells were then washed with an Amplification Enhancer Wash Buffer for 1 minute at room temperature.

The cells were washed with 70% ethanol and followed by 100% ethanol. Autofluorescence quenching was performed by adding the Autofluorescence Solution and incubating for 10 minutes at room temperature in the dark. Cells were then washed with PBS-T, and nuclei were stained using the Xenium Nuclei Staining Buffer. Cells were washed three times with PBS-T and kept in PBS-T until further analysis.

### Custom Probe Design for CRISPR sgRNA Detection

We designed sgRNA detection probes using a 10X Genomics Xenium 480 Gene Custom Panel. We chose this option instead of adding custom probes to the Xenium Prime 5K panel because this is limited to only 100 custom probes. We then designed probes to maximize the amount of unique sequence per sgRNA, with one probe arm overlapping the protospacer and part of the flanking sequence and the other arm overlapping the constant region of the sgRNA. We also optimized the dinucleotides at the ligation junction, avoiding the disfavored dinucleotides CG, CT, GG, and GC. Finally, we aimed for a per-arm melting point between 50°C and 70°C.

### Post-Xenium Immunofluorescence staining and imaging

Post-Xenium slides were stored in PBS with 0.05% Tween (PBS-T) at 4°C until ready for immunofluorescence staining (IF). PBS-T was removed, and slides were washed 3 times with 500 μl of PBS-T and incubated in 500 μl Blocking buffer (1X PBS pH 7.4, 0.1% TWEEN-20, and 10% donkey serum) for one hour at room temperature. The blocking buffer was removed, and 500 μl of primary antibodies for MAP2 (R&D Systems, MAB8304, 1:1000) and NF-H (Abcam, ab4680, 1:50,000) in PBS-T were added. The primary antibodies were incubated overnight at 4°C. Slides were washed 3 times with 500 μl PBS-T before incubation with donkey secondary antibodies (Invitrogen, AlexaFluor, 1:500) for 1 hour at room temperature, followed by 3 more washes. Slides were stored in PBS-T until imaged at 4°C. The post-Xenium IF staining images were acquired using the CellVoyager CQ1 and stitched together using Fiji.

### Gene expression preprocessing, unsupervised clustering and heatmap generation

We performed cell clustering through Scanpy’s single cell analysis pipeline, followed by quality control through the calculate_qc_metrics() function, and cells with high mitochondrial and ribosomal genes were removed. We then did normalization by conducting count depth scaling with subsequent log plus one (log1p) transformation through scanpy’s <u>sc.pp</u>.normalize_total() and <u>sc.pp</u>.log1p() functions. We conducted feature selection by first removing any cells with fewer than 800 transcripts, which left us with 77,455 cells for our dual SMAD neurons and 59,466 cells for our perturbed iNeurons, followed by dimensionality reduction through principal component analysis. Through the <u>sc.pp</u>.neighbors(n_neighbors =50, n_pcs=5) and <u>sc.tl</u>.umap() functions we did neighborhood graph construction with PCA representation of the matrix, followed by visualization and embedding in two dimensions with Uniform Manifold Approximation and Projection (UMAP). Using the <u>sc.pl</u>.umap(color=[“leiden”]) function with a resolution of 2, we clustered through the Leiden graph method. In order to classify each cluster, we conducted differentially-expressed gene analysis through the <u>sc.tl</u>.rank_genes_groups() function and Wilcoxon statistical analysis. We then analyzed the top 30 DEGs from each cluster to identify the cell type classification, with an additional validation step of mapping distinct cell type markers onto our UMAP. For our iNeuron perturbation heatmap we ran an omnibus Kruskal-Wallis test across all gRNA perturbed groups including control without assuming normality, getting one p-value per gene. We then corrected those p-values across genes using Benjamini-Hochberg FDR (alpha < 0.05) under each method and z-scored the per-perturbation log fold changes for significant genes and plotted the clustered heatmap.

### RNA Velocity Analysis

RNA velocity characterizes the rate of change in gene expression at the single-cell level by comparing pre-mature and mature RNA abundances to infer cellular dynamics. Conventionally, this is accomplished by quantifying unspliced (intronic) and spliced (exonic) transcripts, thereby predicting the trajectory of cell differentiation processes. Our Xenium probes do not distinguish between spliced and unspliced transcripts of the same gene. However, our data do provide transcript localization information, allowing us to distinguish between nuclear and cytoplasmic RNA counts for each gene. This can be interpreted as modifying the RNA velocity model to estimate nuclear export rates instead of RNA splicing rates. We quantified nuclear and cytoplasmic transcript counts using the Spaceranger outputs. We then selected the top 2,000 most highly variable genes and performed RNA velocity analysis using our previously published VeloVAE package, employing the full variational Bayes model with default parameters.

### GCaMP signal intensity processing and LASSO regression

At each time point, GCaMP images are acquired in 136 frames (17×8), each frame measuring 2000×2000 pixels with 1% overlap. Prior to stitching, the GCaMP signals undergo median subtraction on a per-pixel basis. The individual frames are then stitched together by calculating signal intensities in the overlapped regions through median-based calibration. Subsequently, the stitched GCaMP image is registered to the Xenium image using 130 aligned reference points.

For each time point, the neural activity of individual cells is computed by averaging GCaMP intensities (within an affine-transformed coordinate space) across the pixel area corresponding to each cell boundary as identified by the Xenium platform.

For each cell, peaks are identified according to three criteria: (1) intensity values that exceed 1.5× the first quartile for ≥2 consecutive time points, (2) intensity values that exceed 2× the minimum for ≥2 consecutive time points, or (3) intensity values above 1.2× the first quartile for ≥3 consecutive time points. Additionally, an absolute difference of 5 is enforced to mitigate false positives in cells with consistently low GCaMP signals. The peak amplitude is measured as the difference between the peak’s maximum and the cell’s first quartile, while the fold amplitude is the ratio of the peak’s maximum to that quartile. Peak duration is calculated by an average of the intervals between the start and end of each peak. The area under the curve (AUC) is obtained by summing all signal intensities above the median across the relevant time points.

In the subsequent LASSO regression analysis, the log1p transformation of the AUC serves as the dependent variable, and only genes expressed in ≥5% of cells are included. The optimal number of variables is selected via cross-validation, ultimately yielding 26 genes for downstream Gene Ontology (GO) enrichment analysis under biological process terms.

### GCaMP-based cell type clustering

Inactive cells are defined as cells with no peak detected. The active cells are clustered using the z-score transformed values of 5 GCAMP features: peak counts, median, fold peak amplitude, peak amplitude, and peak duration, using k-means clustering. Different k-means clustering results are compared, and the 5-clustered result shows the best ability to capture the neural activity heterogeneities.

### Identifying neurite- and filament-enriched transcripts

The immunofluorescence (IF) staining images were first registered with Xenium images to ensure accurate alignment. Subsequently, cell bodies were identified and excluded from the IF images. A threshold was then established for the IF pixel intensity, and after eliminating the overlapping pixels between groups, the remaining pixels were presumed to represent dendritic or axonal structures. The spatial coordinates of these residual pixels were matched with those of the transcripts to determine their localization within dendrites or axons. Finally, differential RNA expression analysis was performed by comparing the expression profiles of the dendritic/axonal regions with those of the cell body regions. UTR lengths were computed using the 5′ and 3′ UTR measurements of the canonical transcript for each gene

### Analysis of BFP intensity and lysosome movement

At each time point, lysosome images are acquired in 480 frames (32×15), each frame measuring 2000×2000 pixels with 2% overlap. We also captured the bright field (BF) and blue fluorescent protein (BFP) channels for each FOV. We stitched the BF channel images of all the FOVs, then registered it with the Xenium image. The BFP intensity of a cell is computed by averaging BFP intensities (within an affine-transformed coordinate space) across the pixel area corresponding to each cell boundary as identified by the Xenium platform. We used Youden’s J statistic to calculate the best threshold to separate cells with and without gRNA expression using BFP intensity, and mark the cells as BFP+ or BFP- using the threshold.

We used the FIJI plugin TrackMate to track the lysosome movement in each FOV, allowing at most 2 frame gaps^32^. The output of TrackMate has the locations of each detected lysosome object and the statistics of lysosome tracks, for example, total distance traveled and mean directional change rate. The lysosome tracks that have at least 2 spots inside and at least 2 spots outside of the soma mask for a cell are assigned to the corresponding cell. Then, we ran statistical analysis to compare each gene perturbation group to the negative control group for each metric of the assigned lysosome tracks. We plotted Kymographs of the lysosome tracks for the gene perturbation group that shows significant metrics compared to negative control. The lysosome tracks are randomly selected, with the restriction that their values of the significant metric should be within a particular percentile range (same percentile range used for negative control and perturbed cell groups). The x-axis of the graph is the closest distance between the detected lysosome object and the cell boundary, and the y-axis is the time point.

### Cellpose-SAM Segmentation Model Training

We conducted automated soma (cell body) and neurite segmentation on the brightfield tiles using the Cellpose-SAM deep learning framework (Cellpose v4.0.5)^39^. Separate custom models were trained for somas and neurites to capture their distinct morphologies. For the soma model, we manually annotated 50 representative tiles (out of 480) and trained the model for 100 epochs (using n_train = 50 images) with a learning rate of 1×10^−5 and weight decay = 0.1. For the neurite model, a smaller training set of 6 annotated tiles was used (n_train = 6, 100 epochs) with a learning rate of 6×10^−5 and weight decay = 0.1. During model training and inference, we applied mild Gaussian smoothing to the input images (Cellpose smooth_radius = 1.0) to reduce noise, consistent with recommended practices (smooth radius ∼1/10 of object diameter). The trained models were then applied to each brightfield tile to generate binary and uint16 mask images for somas and neurites, respectively. Each mask image contains labeled pixels for either soma or neurite objects, and these served as the basis for downstream CellProfiler phenotypic quantification.

### Mask Post-processing and CellProfiler Analysis

Cellpose-generated masks were post-processed and quantified using CellProfiler (v4.2.8)^40^. For each tile, the soma and neurite mask images were imported as objects in CellProfiler via the *ConvertImageToObjects* module. Because the raw neurite masks tended to be fragmented, we performed a morphological closing operation in CellProfiler to connect discontinuous neurite segments. In practice, this was achieved by dilating each neurite object by a radius of ∼7 pixels and merging overlapping fragments, effectively “closing” small gaps. Next, we associated neurites with their parent somas using a seeded propagation approach. Specifically, we ran *IdentifySecondaryObjects* in Propagation mode, treating each soma object as a seed and using the brightfield intensity image as the guidance for neurite growth. This propagation used an adaptive Otsu threshold (two-class) to delineate cell boundaries, allowing neurite regions to expand from each soma until intensity boundaries were reached (tolerating small gaps in the process). The thresholding parameters were tuned as follows: a Gaussian smoothing filter σ≈1.3844 applied to the intensity image, intensity log-transform enabled, adaptive window radius = 450 pixels, lower/upper threshold bounds = 0.0/1.0 (no manual clipping), and regularization λ = 0.05 for the propagation algorithm. These settings ensured that faint neurite extensions attached to a soma were included in that cell’s territory while limiting over-expansion into the background. As a result, we obtained composite cell objects that encompassed each soma and its connected neurite network (denoted as SomaAndNeuritesMerged in our pipeline).

### Quantification of Soma and Neurite Morphometric Features

We compiled a comprehensive set of morphological and intensity features for the segmented objects using CellProfiler modules^40^. First, neurite structures were skeletonized to facilitate length and branch measurements (CellProfiler *ExpandOrShrinkObjects* set to “skeletonize”). We then measured neurite outgrowth characteristics per cell using *MeasureObjectSkeleton* (recording total neurite length, branch counts, endpoints, etc.). Basic morphometry of each object was captured with *MeasureObjectSizeShape* (e.g. area, perimeter, shape descriptors).

Spatial relationships were quantified: *MeasureObjectNeighbors* counted proximal relationships (e.g. number of neurite segments touching each soma), and *MeasureObjectOverlap* calculated the overlap and minimum distances between soma and neurite objects (including an earth mover’s distance calculation based on object skeletons). We also assessed texture and granularity of staining within each object: *MeasureTexture* computed Haralick texture features for each soma and neurite using the brightfield and fluorescence images, while *MeasureGranularity* evaluated the presence of intensity patterns at multiple spatial scales (using structuring element radii up to 10 px). In addition, global image-level metrics were obtained.

*MeasureImageSkeleton* was applied to the neurite mask as a whole to gauge network connectivity across each field (e.g. total neurite length per image and number of branch points). *MeasureImageIntensity* and *MeasureImageQuality* were used to evaluate overall image intensity distribution (within and outside objects) and image focus/quality metrics, respectively. We also calculated the fractional area occupied by cellular material in each tile using *MeasureImageAreaOccupied*, focusing on soma, neurite, and combined soma+neurite object masks. All measured features were exported via CellProfiler’s spreadsheet export module. Data were organized by object category, including experiment-level and image-level measurements, as well as per-object measurements for each identified object set: soma, neurite, SkeletonNeurite (individual neurite skeleton segments), SkeletonMergedNeurite (skeletons after neurite merging), and SomaAndNeuritesMerged (the combined soma-with-neurites objects).

### Analysis of cell morphology from Cellpose-SAM and CellProfiler results

After we got the merged soma and neurite masks from Cellpose-SAM, we matched the pixels in the cell masks with the transcript locations that we obtained from the Xenium experiment. We assigned the Cellpose-SAM-segmented cells to different gene perturbation groups based on the expression of gRNAs, and compared each gene perturbation group with the negative control group for the cell morphology metrics that we obtained from CellProfiler. We displayed the outlines of the cell masks on the BF channel images to show the cell morphology.

## Acknowledgements.

This work was supported by NIH grant R01HG010883 to J.D.W., University of Michigan Research Scout grant OORRS033123 to J.D.W., and T32GM141746 to N.Z.

## Data Availability

Raw and processed data are available on GEO: GSE304064.

---

**Supplementary Figure 1:**
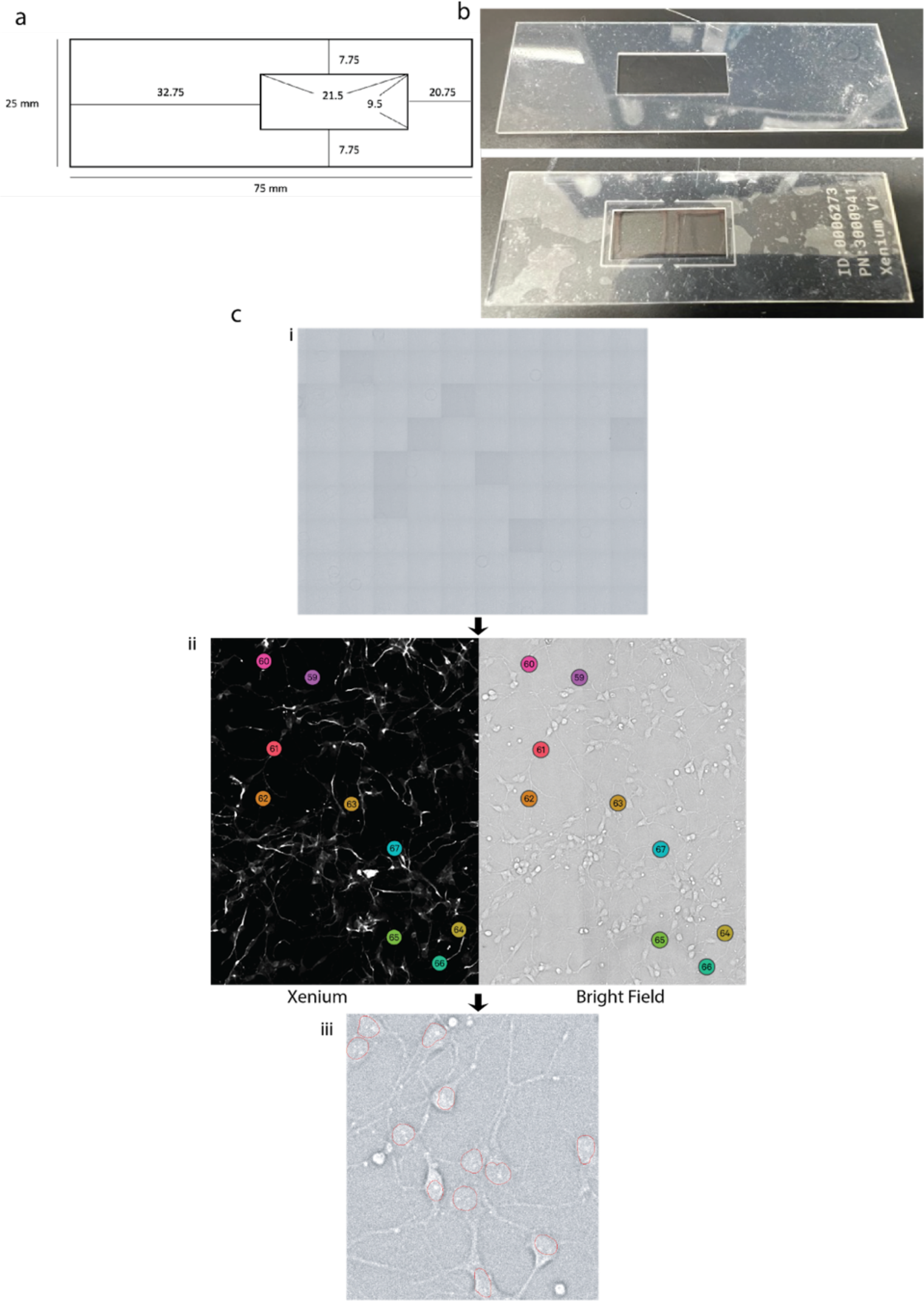
Gasket design and Xenium registration **a**. Polydimethylsiloxane (PDMS) gasket design for Xenium slides. **b**. Image of a PDMS gasket (top). Gasket placed on Xenium slide (bottom). **c**. Workflow of registration and confirmation. **(i)** All fields of view for pre-Xenium brightfield images were stitched together. **(ii)** A screenshot of Xenium (left) and stitched brightfield image (right) alignment. Key points (circled with a number) were placed on each image to register the pre-Xenium images with the Xenium images, allowing us to identify the same cells in both images. **(iii)** Xenium segmentation masks shown on pre-Xenium images, confirming registration quality. The red dotted circles (cell body from Xenium output) are registered to the cell bodies from the brightfield image.

**Supplementary Figure 2:**
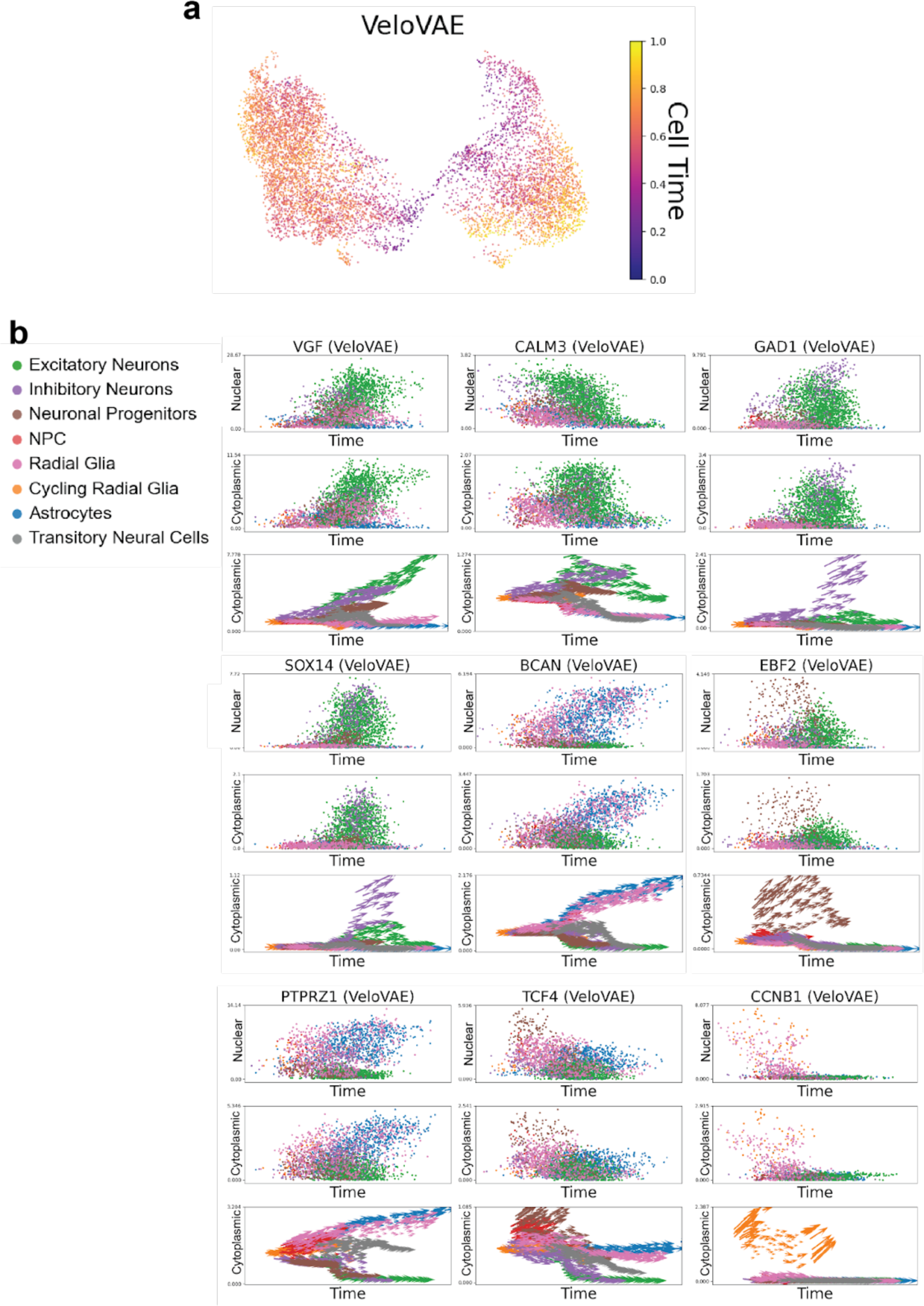
RNA velocity latent time and gene-level cell dynamics **a.** UMAP coordinates colored by latent time from VeloVAE result. **b**. Observed values for nuclear RNA, cytoplasmic RNA, and RNA velocity plotted as a function of latent time and colored by cell types for 9 different genes.

**Supplementary Figure 3:**
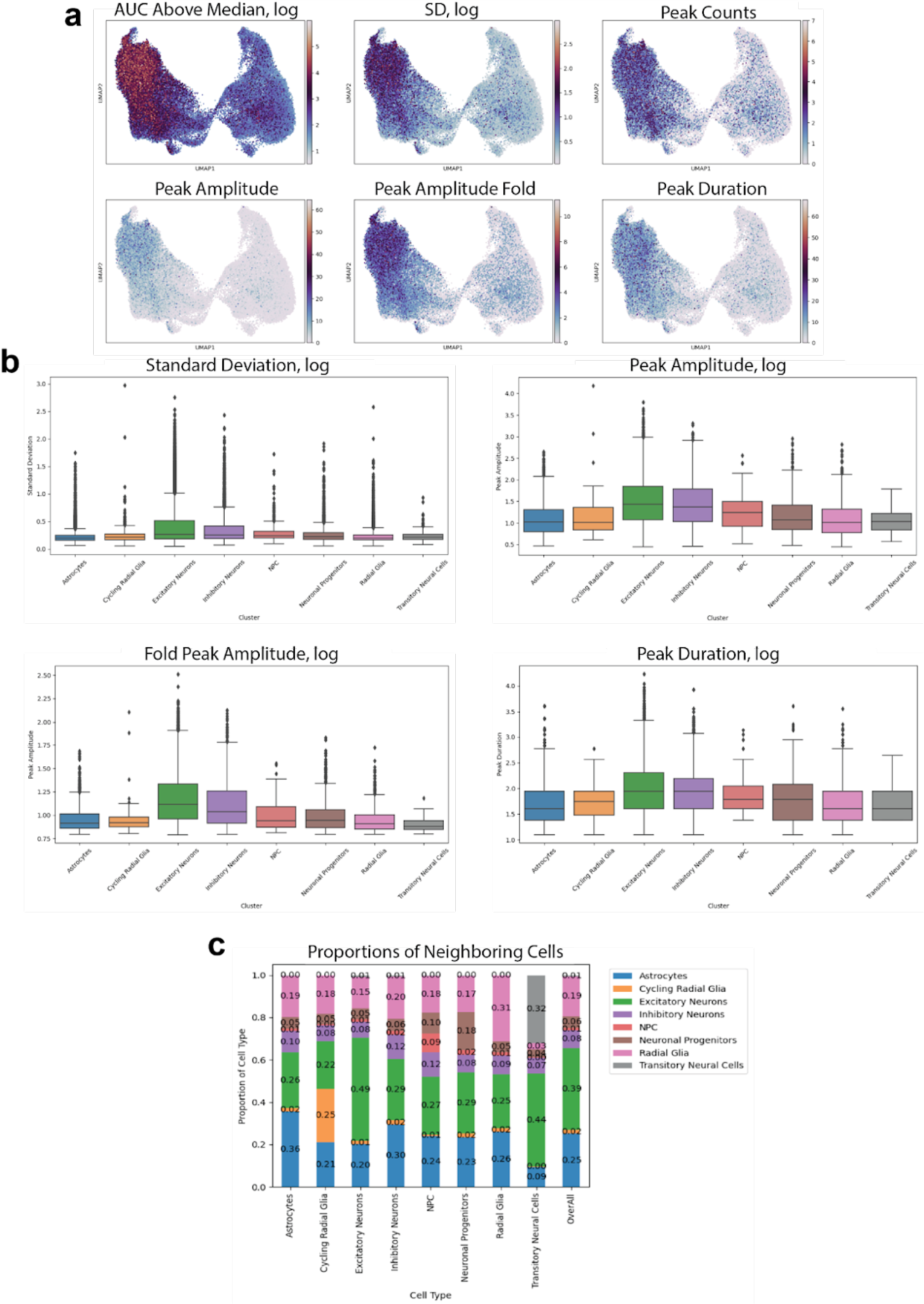
Supplementary GCaMP metrics and neighboring cell type proportions **a.** UMAP coordinates colored by GCaMP metrics, including AUC above median (log transformed), standard deviation (log transformed), peak counts, peak amplitude, peak amplitude fold change, and peak duration. **b**. Box plots of log-transformed standard deviation, peak amplitude, peak amplitude fold change, and peak duration for different transcriptomic clusters. **c**. Proportions of spatially neighboring transcriptomic clusters. Cells that are clustered in the same cell type tend to be more likely to occur near each other, especially for the rare cell clusters (cycling radial glia and transitory neural cells). Neighboring cells are defined as cells whose nearest distance (calculated between soma masks) is closer than 2.125 microns.

**Supplementary Figure 4:**
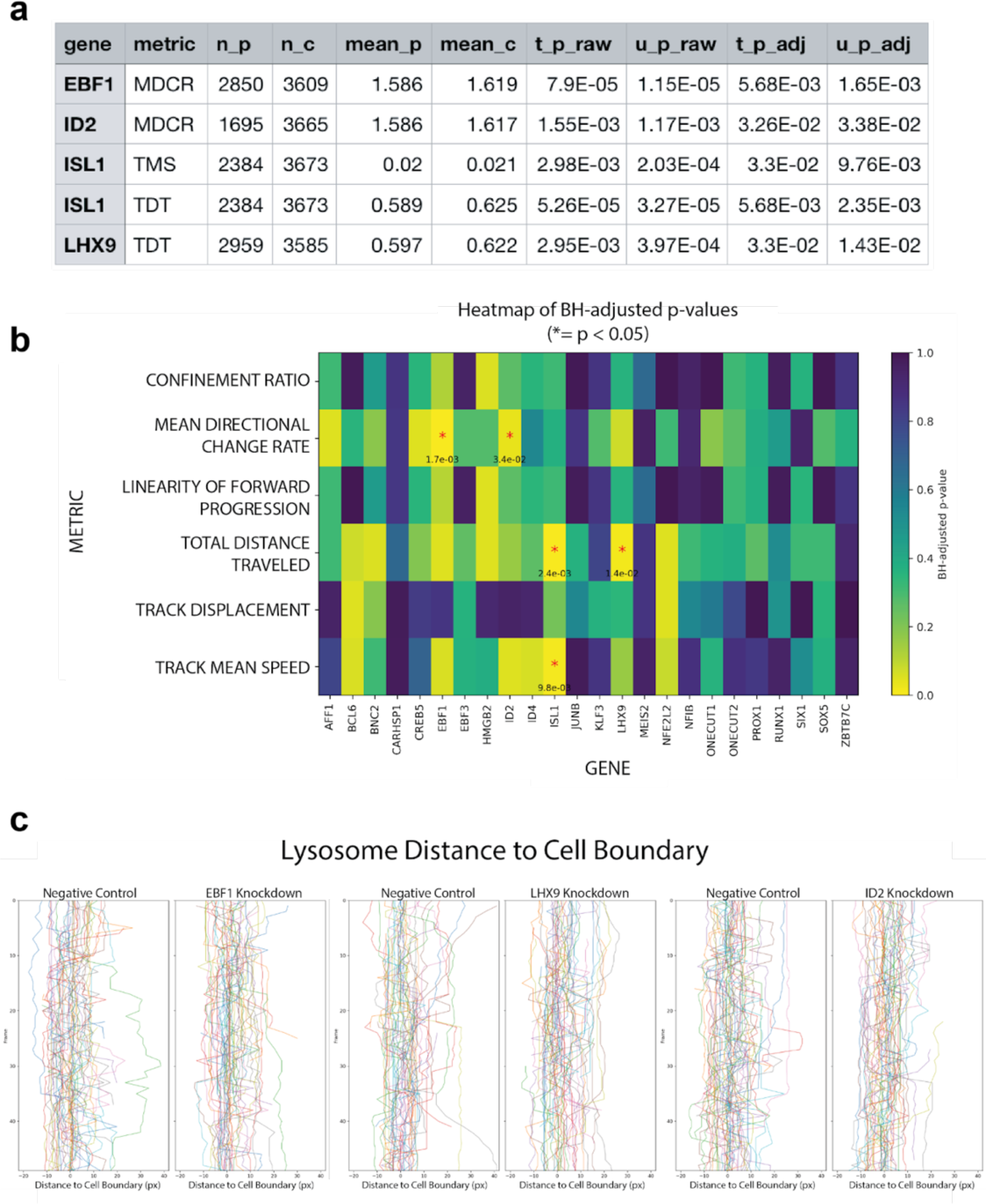
Additional lysosome analysis **a.** Table of the gene metric pairs that have BH-adjusted Mann-Whitney U test p-values < 0.05. n_p (number of perturbation group cells), n_c (number of negative control group cells), mean_p (mean value of perturbation group cells), mean_c (mean value of negative control group cells), t_p_raw (raw p-value of t-test), u_p_raw (raw p-value of Mann-Whitney U test), t_p_adj (BH adjusted p-value of t-test), u_p_adj (BH adjusted p-value of Mann-Whitney U test), MDCR (mean direction change rate), TMS (track mean speed), TDT (total distance traveled). **b.** Heatmap of BH-adjusted p values. Significant gene metric pairs are marked by asterisks. **c.** Kymographs of lysosome distance to cell boundary for EBF1, LHX9, and ID2 KD groups compared to the negative control. Lysosomes were randomly selected from matched percentiles of the corresponding lysosome metric in each group to ensure representative sampling.

**Supplementary Figure 5:**
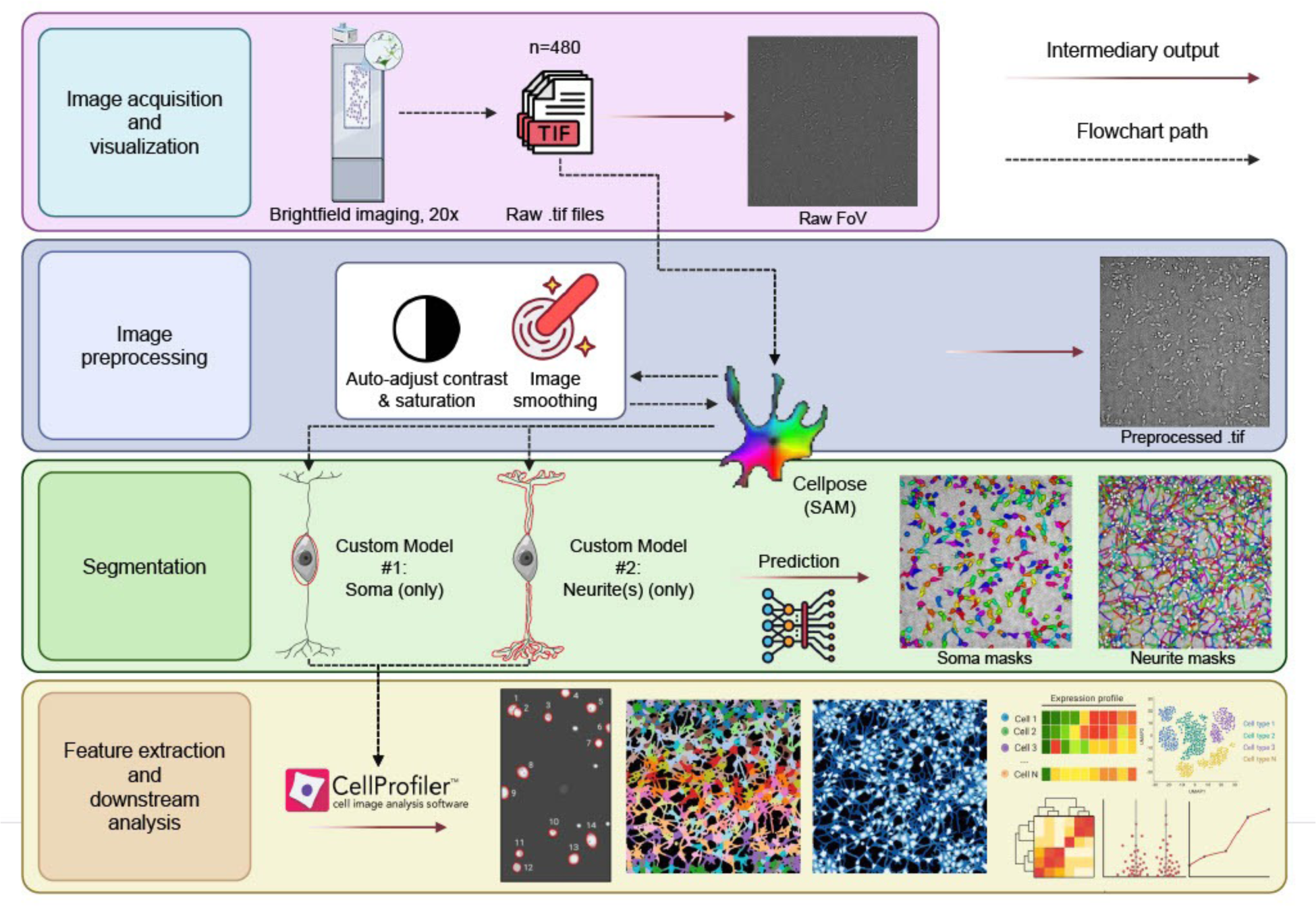
Overview schematic of the post-xenium high-content image processing workflow for human iPSC-derived neuron cultures. Brightfield images (20× magnification; Yokogawa CQ1 system) were acquired as 480 tiled fields of view and underwent Cellpose (v4.0.5) image filtering preprocessing (e.g., auto contrast enhancement and smoothing) to enhance downstream segmentation. The pipeline branches into two separate custom segmentation models based on the Cellpose-SAM architecture (Cellpose v4.0.5, Segment Anything Model): one model specialized for identifying neuronal somas (cell bodies) and another for tracing neurites (thin axonal and dendritic processes). Example output masks from each model are depicted, highlighting isolation of somas vs. neurites from the same input image. In the final step, segmented objects are analyzed with CellProfiler (v4.2.8) to extract quantitative, object-level morphometric and phenotypic features. This includes overlaying identified objects on the original images, generating heatmaps of feature distributions, and calculating inferred metrics (e.g., size, shape, and intensity) for each segmented neuron and its associated neurites.

**Supplementary Figure 6:**
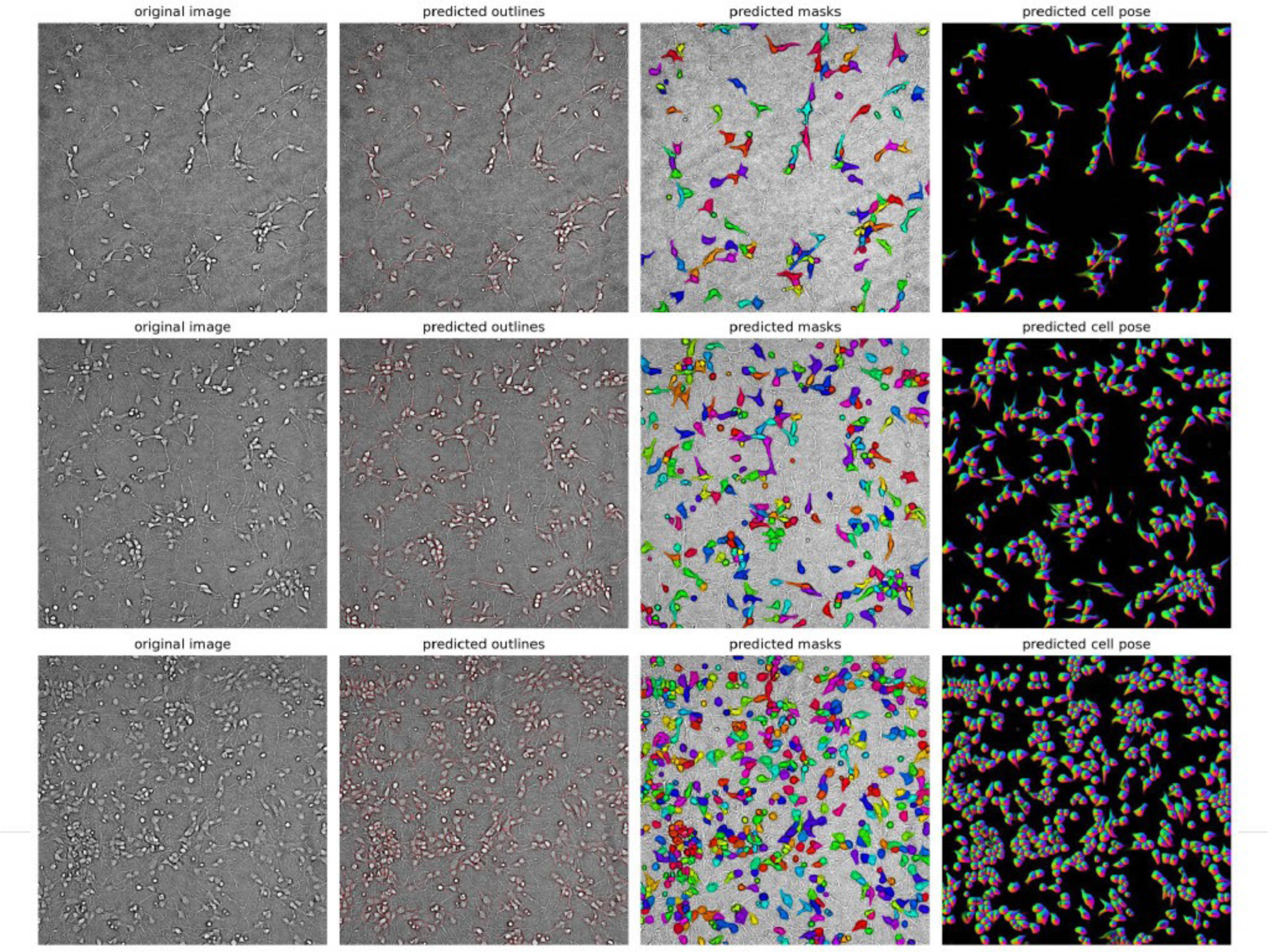
Examples of post-xenium soma segmentation results. As in Supplementary Figure 2, Representative brightfield segmentation results from neuronal somas obtained using a custom Cellpose-SAM model (Cellpose v4.0.5). Each row illustrates a distinct field of view (FOV) from the 20× brightfield CQ1 dataset, showing segmentation performance across different cell densities (top to bottom: sparse, intermediate, and high). For each FOV, panels show (left to right): the original preprocessed brightfield image, the Cellpose-predicted soma boundaries (outlined in cyan), the color-coded segmentation masks overlaid onto the original image indicating the identified individual soma objects, and the model’s internal cell pose rendering (spatial flow-field visualization) depicting the directionality and extent of each segmented soma.

**Supplementary Figure 7:**
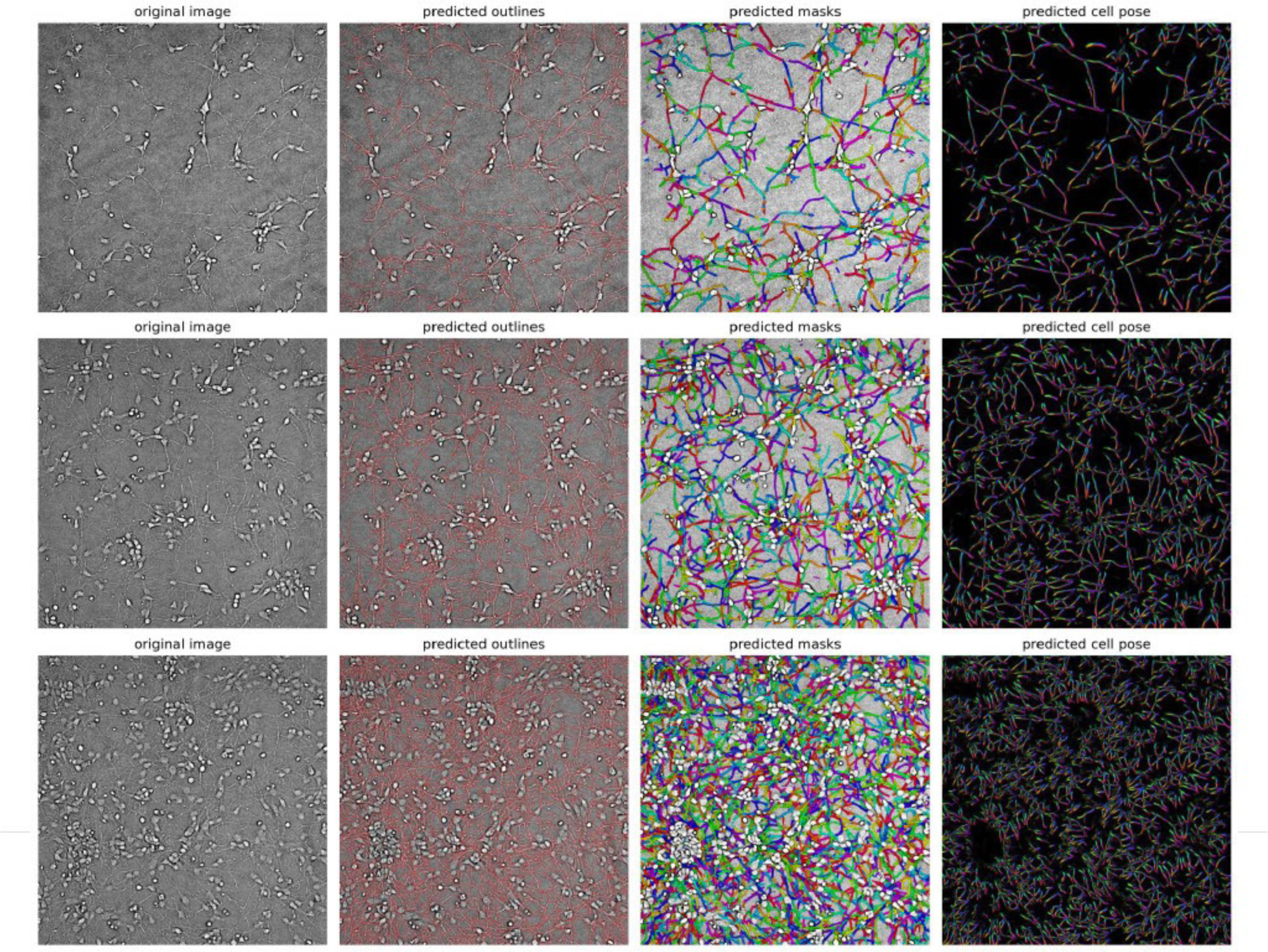
Examples of post-xenium neurite segmentation results. As in Supplementary Figure 2, representative brightfield segmentation results from neuronal neurites obtained using a custom Cellpose-SAM model (Cellpose v4.0.5). Each row illustrates a distinct field of view (FOV) from the 20× brightfield CQ1 dataset, capturing segmentation of fine neurite structures across different cell densities (top to bottom: sparse, intermediate, and high). For each FOV, panels show (left to right): the original preprocessed brightfield image, the Cellpose-predicted neurite boundaries (outlined in cyan, tracing thin filamentous processes), the color-coded segmentation masks overlaid onto the original image indicating individual neurite segments, and the model’s internal cell pose rendering (spatial flow-field visualization) depicting the continuity, directionality, and extent of each segmented neurite.

**Supplementary Figure 8:**
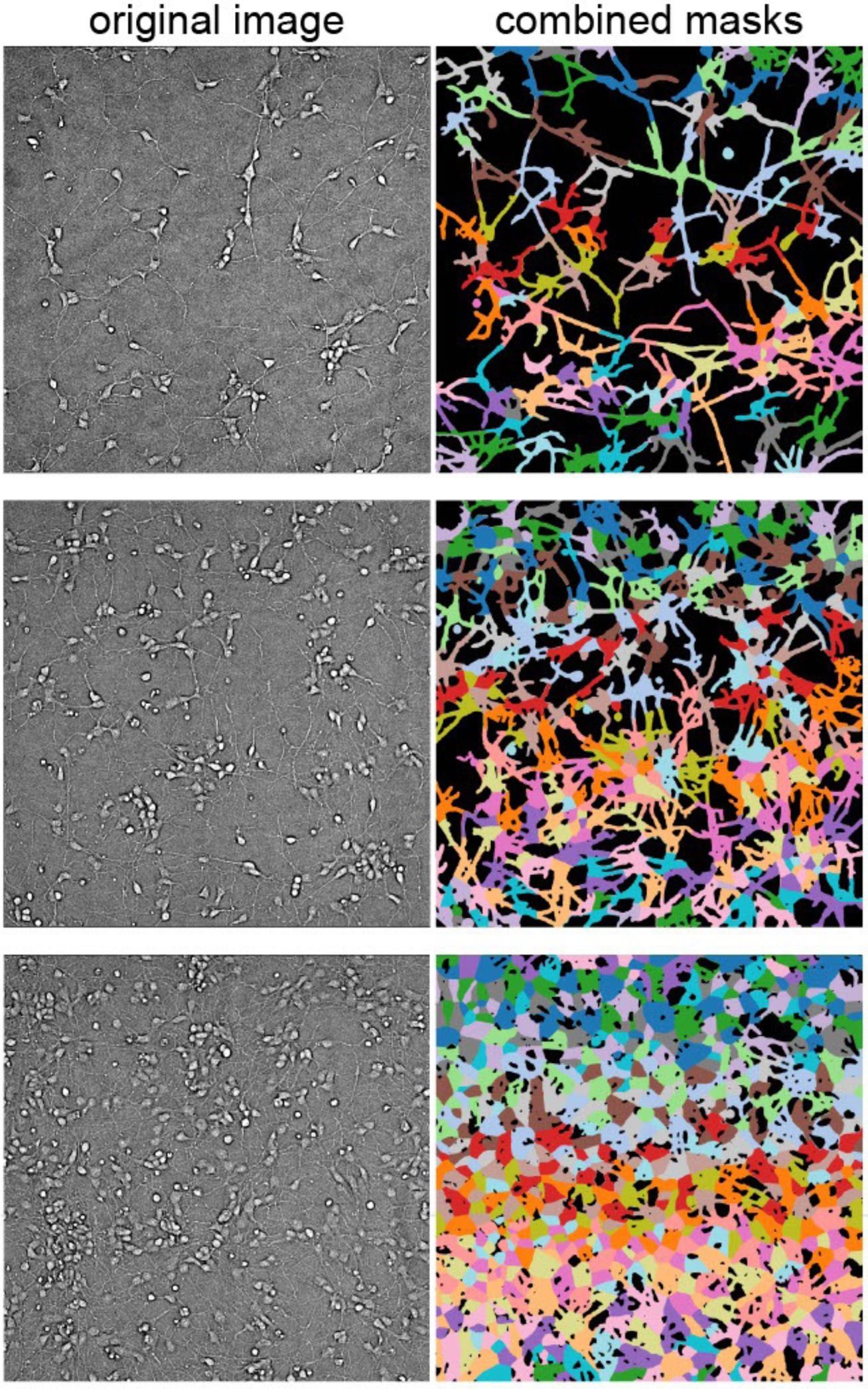
Combined soma-neurite object segmentation. Representative results illustrating the merging of soma and neurite segmentation masks into contiguous combined neuronal objects using CellProfiler (v4.2.8). Each row displays a distinct field of view (FOV) from the 20× brightfield CQ1 dataset across different cell densities (top to bottom: sparse, intermediate, and high). For each FOV, panels show the original brightfield image (left) alongside the corresponding combined masks (right), where each uniquely colored object represents a soma and its associated neurite extensions after morphological closing and propagation-based mask merging. Parameters for mask merging included adaptive Otsu thresholding, morphological closing (disk radius = 7 pixels), adaptive window size (450 pixels), and propagation regularization (λ = 0.05).

**Supplementary Figure 9:**
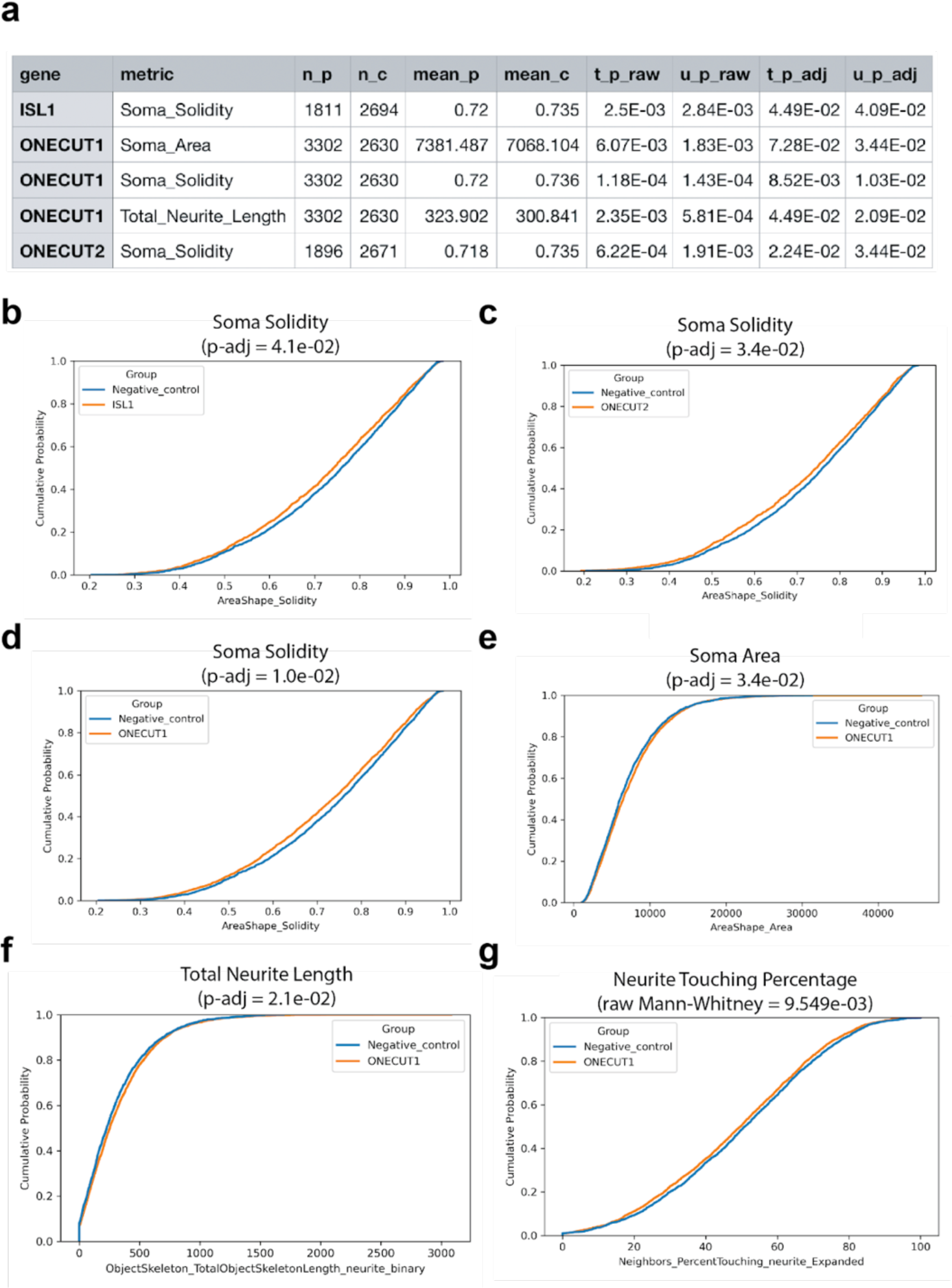
Additional cell morphology analysis **a.** Table of the gene metric pairs that have BH-adjusted Mann-Whitney U test p-values < 0.05. Soma solidity measures the proportion of the pixels in the convex hull that are also in the object, i.e., ObjectArea/ConvexHullArea. A smaller number indicates the object shape is more irregular. n_p (number of perturbation group cells), n_c (number of negative control group cells), mean_p (mean value of perturbation group cells), mean_c (mean value of negative control group cells), t_p_raw (raw p-value of t-test), u_p_raw (raw p-value of Mann-Whitney U test), t_p_adj (BH adjusted p-value of t-test), u_p_adj (BH adjusted p-value of Mann-Whitney U test). **b**. CDF plot of ISL1 KD cells compared to the negative control for soma solidity. **c**. CDF plot of ONECUT2 KD cells compared to the negative control for soma solidity. **d**. CDF plot of ONECUT1 KD cells compared to the negative control for soma solidity. **e**. CDF plot of ONECUT1 KD cells compared to the negative control group for soma area. **f**. CDF plot of ONECUT1 KD cells compared to the negative control for total neurite length. **g**. CDF plot of ONECUT1 KD cells compared to the negative control for neurite touching percentage. The metric is not significant after BH correction. However, the value is lower for the ONECUT1 KD group compared to the negative control.

